# Serial horizontal transfer of vitamin-biosynthetic genes enables the establishment of new nutritional symbionts in aphids’ di-symbiotic systems

**DOI:** 10.1101/556274

**Authors:** Alejandro Manzano-Marín, Armelle Coeur d’acier, Anne-Laure Clamens, Céline Orvain, Corinne Cruaud, Valérie Barbe, Emmanuelle Jousselin

**Affiliations:** UMR 1062 Centre de Biologie pour la Gestion des Populations, INRA, CIRAD, IRD, Montpellier SupAgro, Univ. Montpellier, Montpellier, France; Institut de Biologie François-Jacob, CEA, Genoscope, Évry Cedex, France

**Keywords:** symbiosis, mutualism, horizontal gene transfer, vitamin biosynthesis, *Erwinia haradaeae*

## Abstract

Many insects depend on obligate mutualistic bacteria for the provisioning of essential nutrients lacking from their food source. Most aphids (Hemiptera: Aphididae), whose diet consists of phloem, rely on the bacterial endosymbiont *Buchnera* for the supply of essential amino acids and B vitamins. However, in some aphid lineages provision of these nutrients is partitioned between *Buchnera* and an extra bacterial partner. Little is known about the origin and the evolutionary stability of these di-symbiotic systems. Here, we explore these questions in a group of aphids which harbours both *Buchnera* and an *Erwinia*-related symbiont. Using fluorescence *in situ* hybridisation, we located the *Erwinia* symbiont in close proximity to *Buchnera* where it inhabits its own bacteriocytes. Analyses of whole-genome sequences of the endosymbionts of 9 aphid species show that *Erwinia* genomes are highly reduced, syntenic, and display phylogenetic congruency with *Buchnera*. Altogether these results depict a scenario for *Erwinia* that mirrors the evolutionary history of *Buchnera*: its intracellularisation as an obligate partner is a one time event that has resulted in drastic genome reduction and long-term co-divergence with its host. Additionally, we found that the *Erwinia* genes that complement *Buchnera*’s deficiencies, but also a set of genes that carry a new nutritional function, have actually been horizontally acquired from a *Sodalis*-related bacterium. A subset of these genes have been further transferred to a new *Hamiltonella* co-obligate symbiont in a specific *Cinara* lineage. This shows that the establishment and dynamics of multi-partner endosymbioses can be mediated by lateral gene transfers between co-ocurring symbionts.

## Introduction

Obligate symbiotic bacteria are pervasive among different groups of animals whose diets lack essential nutrients. These include blood-(Manzano-Marín *et al.*, 2015; Smith *et al.*, 2015; Říhová *et al.*, 2017) and sap-feeders (xylem and phloem) (Husnik and McCutcheon, 2016; Matsuura *et al.*, 2018; Santos-Garcia *et al.*, 2014). Aphids (Hemiptera: Aphididae) are plant-sucking insects whose diet consists of plant phloem, which is poor in essential amino acids and B vitamins (Akman Gündü z and Douglas, 2009; Sandstrom and Moran, 1999; Ziegler, 1975). Accordingly, they depend on obligate symbiotic bacteria from the *Buchnera* genus to supply these nutrients. *Buchnera* localises to the bacteriome (Buchner, 1953), a specialised organ made up mainly of cells called bacteriocytes which house the *Buchnera* bacteria intracellularly (Griffiths and Beck, 1975; Munson *et al.*, 1991). Similarly to other “long-term” vertically-inherited obligate endosymbionts, *Buchnera* strains have highly reduced genomes, show a high degree of genome synteny (Tamas *et al.*, 2002; van Ham *et al.*, 2003), and display strong phylogenetic congruency with their hosts (Baumann *et al.*, 1995; Funk *et al.*, 2000; Jousselin *et al.*, 2009); evidencing an early drastic genome reduction followed by co-divergence with their hosts.

Aphids can also harbour additional facultative endosymbionts, which are neither required for normal development nor are generally fixed across populations of the species (Oliver *et al.*, 2010). These secondary bacterial associates can have beneficial effects for their hosts, such as protection against parasitoids (Hansen *et al.*, 2012; Oliver *et al.*, 2003) or survival after heat stress (Chen *et al.*, 2000; Montllor *et al.*, 2002). Some aphids have established new obligate nutritional-based associations with additional bacterial lineages. The best known cases occur in species from the Lachninae subfamily of aphids, where *Buchnera* symbionts have ancestrally lost the capability to synthesise two essential B vitamins: biotin (B_7_) and riboflavin (B_2_) (Manzano-Marín *et al.*, 2016; Meseguer *et al.*, 2017). As a consequence, both the aphids and their *Buchnera* now rely on fixed secondary endosymbionts to complement the nutritional deficiencies. These younger co-obligate symbionts belong to diverse gammaproteobacterial taxa known to be facultative endosymbionts of aphids: *Sodalis*, *Fukatsuia*, and *Serratia symbiotica* (Manzano-Marín *et al.*, 2017; Meseguer *et al.*, 2017). Contrary to *Buchnera*, these symbionts present more diverse cell shapes and tissue tropism. They can be round or rod-shaped confined to the cytoplasm of bacteriocytes, or they can display long filamentous cells that are found surrounding bacteriocytes, co-infecting *Buchnera*’s bacteriocytes, or inhabiting their own (Buchner, 1953; Fukatsu and Ishikawa, 1993; Manzano-Marín *et al.*, 2017; Pyka-Fościak and Szklarzewicz, 2008).

In *Cinara* aphids (the most species-rich genus within the Lachninae), several cases of secondary co-obligate symbiont acquisitions and replacements have been proposed (Manzano-Marín *et al.*, 2017; Meseguer *et al.*, 2017). This turnover of the newly arrived companion symbionts is actually often observed in multi-partner endosymbioses (Husnik and McCutcheon, 2016). An ancestral reconstruction of the bacterial symbiotic associations within this aphid clade suggested that the common ancestor of these aphids harboured a *Se. symbiotica* endosymbiont which has been subsequently replaced several times by other bacterial taxa (Meseguer *et al.*, 2017). These include the aforementioned *Sodalis* and *Fukatsuia*, as well as the gammaproteobacteria *Erwinia*. While the formers have closely related relatives found living as facultative or recently acquired co-obligate symbionts in aphid species (Guay *et al.*, 2009; Heyworth and Ferrari, 2015; Manzano-Marín *et al.*, 2017; Meseguer *et al.*, 2017), *Erwinia*, to our knowledge, has only been identified as part of the gut flora of *Acyrthosiphon pisum*.

*Erwinia* bacteria are primarily known as plant-pathogenic and plant-associated bacteria (Kado, 2006), with genomes sizes ranging between 3.83 and 5.07 Mbps (based on currently available fully closed genomes). Regarding *Erwinia* and aphids, early studies isolated and axenically cultured an aerobic *Erwinia* bacterium from the gut of a laboratory strain of the pea aphid *Ac. pisum*, named *Erwinia aphidicola* (Harada *et al.*, 1997). This bacterial species, along with that termed Y, were the dominant groups, among the seven bacterial ones, which were found to constitute the major flora of the aphid’s gut (Harada and Ishikawa, 1993; Harada *et al.*, 1996). In *Cinara* aphids, *Erwinia* bacteria have been found to be fixed in a particular monophyletic group of *Cinara* aphids (Meseguer *et al.*, 2017). Moreover, the study showed that at least one strain of these *Erwinia* symbionts has a reduced genome with the predicted capability to supplement *Buchnera*’s biotin and riboflavin auxotrophy. Given the above, this association is of particular interest due to **(1)** the possible recent and unusual transition of *Erwinia* from a free-living life-habit to an obligate nutritional endosymbiont and **(2)** its phylogenetically conserved association with its aphid hosts in an otherwise unstable multi-partner endosymbiosis. Its study represents a unique opportunity to understand the mechanisms by which new bacteria establish themselves and persists in multi-partner endosymbioses. In order, to understand the evolutionary history of *Erwinia* as a recently acquired nutritional endosymbiont in aphids; we aimed at answering three sets of questions: **(1)** Are these symbionts derived from a single acquisition of a free-living *Erwinia* lineage and have since cospeciated with their aphid hosts (as shown for the primary symbiont *Buchnera*)? Or alternatively have they been acquired and lost several times, as is probably the case for obligate *Se. symbiotica* in *Cinara* aphids (Manzano-Marín and Latorre, 2016)? **(2)** According to their genomic contents, how do the bacterial partners partition the synthesis of the host’s essential nutrients and does this partitioning change throughout the evolutionary history of the endosymbiosis? For instance, did *Erwinia* bring additional functions to the symbiotic consortia and did it undergo genome reduction upon its integration as an obligate endosymbiont? **(3)** What is the localisation pattern of *Erwinia* within the aphid? Do they inhabit they own bacteriocytes in close proximity with *Buchnera* or do they live in the aphid haemolymph as many facultative endosymbionts? The answers to the latter will inform us on *Erwinia*’s process of integration with the aphid hosts and the possibility of direct interactions between *Buchnera* and *Erwinia*. Altogether these investigations will unravel the factors that are pivotal to the *Buchnera*-*Erwinia* mutualism and its long term stability.

To answer these questions, we first used fluorescence *in situ* hybridisation (**FISH**) microscopy to explore the localisation of *Erwinia* within the aphid. Then, we sequenced, assembled, and annotated the genomes of the co-existing endosymbionts of nine *Erwinia*-associated *Cinara* species. We first conducted phylogenomic analyses to investigate *Buchnera*-*Erwinia* codiversification scenarios and explored the genome-based metabolic complementation displayed by *Buchnera*-*Erwinia* symbiont pairs. These approaches revealed a third co-obligate *Hamiltonella* endosymbiont present in two host species. Through a combination of BLAST searches and phylogenetic analyses, we determined that key genes involved in the symbionts’ nutritional roles have actually been horizontally acquired by *Erwinia* and subsequently passed on to *Hamiltonella*.

## Materials and Methods

### Aphid collection, DNA extraction, and sequencing

Following Meseguer *et al.* (2017), we collected nine different species of *Erwinia*-associated *Cinara* aphid species across the north-western and central USA and the south-east of France (supplementary table S1, Supplementary Material online) and stored in 70% ethanol at 6°C. For whole-genome sequencing, we prepared DNA samples enriched with bacteria following a modified version of the protocol by Charles and Ishikawa (1999) as described in Jousselin *et al.* (2016). For this filtration protocol 15 aphids from a single colony were pooled together. Extracted DNA was used to prepare two custom paired-end libraries in France Génomique. Briefly, 5ng of genomic DNA were sonicated using the E220 Covaris instrument (Covaris, USA). Fragments were end-repaired, 3’-adenylated, and NEXTflex PCR free barcodes adapters (Bioo Scientific, USA) were added by using NEBNext® Ultra II DNA library prep kit for Illumina (New England Biolabs, USA). Ligation products were then purified by Ampure XP (Beckman Coulter, USA) and DNA fragments (*>*200 bp) were PCR-amplified (2 PCR reactions, 12 cycles) using Illumina adapter-specific primers and NEBNext® Ultra II Q5 Master Mix (NEB). After library profile analysis by Agilent 2100 Bioanalyser (Agilent Technologies, USA) and qPCR quantification using the KAPA Library Quantification Kit for Illumina Libraries (Kapa Biosystems, USA), the libraries were sequenced using 251 bp paired-end reads chemistry on a HiSeq2500 Illumina sequencer.

### Fluoresence *in situ* hybridisation microscopy

From the aforementioned nine species of *Erwinia*-associated *Cinara* species, we investigated symbiont localisation patterns in individuals from *Cinara cuneomaculata*, *Cinara kochiana kochiana* (hereafter referred to as *C. kochiana*), and *Cinara curvipes* in modified Carnoy’s fixative (6 chloroform: 3 absolute ethanol: 1 glacial acetic acid) and left overnight, following the protocol of Koga *et al.* (2009). Individuals were then dissected in absolute ethanol to extract embryos and transferred into a 6% solution of H_2_O_2_ diluted in absolute ethanol and were then left in this solution for 2 weeks (changing the solution every 3 days). Then, embryos were washed twice with absolute ethanol. Hybridization was performed overnight at 28°C in standard hybridization buffer (20 mM Tris–HCl [pH 8.0], 0.9 M NaCl, 0.01% SDS, and 30% formamide) and then washed (20 mM Tris–HCl [pH 8.0], 5 mM EDTA, 0.1 M NaCl, and 0.01% SDS) before slide preparation. The embryos of up to 10 individuals were viewed under a ZEISS LSM700 confocal microscope. We used two competitive specific probes for *Buchnera* (**BLach-FITC** [5’-FITC-CCCGTTYGCCGCTCGCCGTCA-FITC-3’], Manzano-Marín *et al.* 2017) and *Erwinia* symbionts in *Cinara* (**ErCinara-Cy3** [5’-ErCinara-Cy3-CCCGTTCGCCACTCGTCGBCA-3’], this study). TIFF-formatted images were exported using the **ZEN** v2.3 SP1 software from ZEISS automatically setting the intensity minimum to Black and the intensity maximum to White value (Auto Min/Max option). Exported images were imported into **Inkscape** v0.92.4 for building the published figures.

### Genome Assembly and Annotation

Illumina reads were right-tail clipped (minimum quality threshold of 20) using **FASTX-Toolkit** v0.0.14 (http://hannonlab.cshl.edu/fastxtoolkit/, last accessed February 7 2019). Reads shorter than 75 were dropped. Additionally, **PRINSEQ** v0.20.4 (Schmieder and Edwards, 2011) was used to remove reads containing undefined nucleotides as well as those left without a pair after the filtering and clipping process. The resulting reads were assembled using **SPAdes** v3.10.1 (Bankevich *et al.*, 2012) with the options --only-assembler option and k-mer sizes of 33, 55, 77, 99, and 127. From the resulting contigs, those that were shorter than 200 bps were dropped. The remaining contigs were binned using results from a **BLASTX** (Altschul, 1997) search (best hit per contig) against a database consisting of the Pea aphid’s proteome and a selection of aphid’s symbiotic bacteria and free-living relatives’ proteomes (supplementary table S2, Supplementary Material online). When no genome was available for a certain lineage, closely related bacteria were used. The assigned contigs were manually screened using the **BLASTX** web server (searching against the nr database) to insure correct assignment. No scaffold was found to belong to a *Sodalis*-related bacterium nor contained a 16S rRNA gene from such bacterial taxa. This binning process confirmed the presence of the previously reported putative *Erwinia* and *Hamiltonella* co-obligate symbionts, as inferred from 16S rRNA gene fragment sequencing from several specimens of the 9 Cinara species analysed in this study (Jousselin *et al.*, 2016; Meseguer *et al.*, 2017), as well as other additional symbionts. For all samples, we identified an additional circular molecule in the *Erwinia* bin representing a putative plasmid. The resulting contigs were then used as reference for read mapping and individual genome assembly using **SPAdes**, as described above, with read error correction.

The resulting *Buchnera* and *Erwinia* genomes were annotated using **Prokka** v1.12 (Seemann, 2014). In order to validate start codons, ribosomal binding sites were predicted using **RBSfinder** (Suzek *et al.*, 2001). This was followed by non-coding RNA prediction using **infernal** v1.1.2 (Nawrocki and Eddy, 2013) (against the **Rfam** v12.3 database (Nawrocki *et al.*, 2015)), **tRNAscan-SE** v1.3.1 (Lowe and Eddy, 1997), and **ARAGORN** v1.2.36 (Laslett, 2004). This annotation was followed by manual curation of the genes on **UGENE** v1.28.1 (Okonechnikov *et al.*, 2012) through on-line **BLASTX** searches of the intergenic regions as well as through **BLASTP** and **DELTA-BLAST** (Boratyn *et al.*, 2012) searches of the predicted ORFs against NCBI’s nr database. Priority for the BLAST searches was as follows: **(1)** against *Escherichia coli* K-12 substrain MG1655 (for both *Buchnera* and *Erwinia*), **(2)** against **Erwinia amylovora** CFBP1430 (for *Erwinia* symbionts), and **(3)** against the whole nr database. For each one of these searches, a match was considered valid following manual inspection which took into account identity, domain match, and synteny. The resulting coding sequences (CDSs) were considered to be putatively functional if all essential domains for the function were found, if a literature search supported the truncated version of the protein as functional in a related organism, or if the CDS displayed truncations but retained identifiable domains (details of the literature captured in the annotation files). For *Hamiltonella* symbionts, we performed a draft annotation using **Prokka** v1.12 and search and curated over 100 selected genes related to essential amino acids, B vitamins, and other co-factors. For *Erwinia* and *Buchnera* symbionts, pseudogenes were also searched based on synteny against closely related available genomes. This prediction was performed using a combination of sequence alignment with **m-coffee** (Wallace, 2006) and **BLASTX** searches against the NCBI’s nr database (restricted to *Erwinia* or *Buchnera* taxon ID). This allowed the identification of pseudogenes by the previous searches. Metabolic reconstruction for Buchnera and Erwinia was performed in **Pathway Tools** v22.5 (Karp *et al.*, 2016) and manually curated using **EcoCyc** (Keseler *et al.*, 2017) and **BioCyc** (Karp *et al.*, 2017). Visual plotting of the inferred metabolism was done by hand using Inkscape v0.92.4.

The annotated genomes of *Buchnera* and *Erwinia*, as well as the unannotated ones of *Hamiltonella* and sequencing reads are available at the European Nucleotide Archive under project numbers PRJEB15506, PRJEB31183, PRJEB31187, PRJEB31188, PRJEB31190, PRJEB31191, PRJEB31194, PRJEB31195, and PRJEB31197.

### Phylogenetic Analyses

For performing both phylogenetic inferences and analysing the genetic differences in *Buchnera* and *Erwinia* from the different aphids, we first ran an orthologous protein clustering analysis using **OrthoMCL** v2.0.9 (Chen *et al.*, 2007; Li, 2003) on two sets of proteins: i) *Buchnera* strains from this study + *Buchnera* from *Cinara cedri* as outgroup, and ii) *Erwinia* + *Pantoea* strains + *Erwinia* endosymbionts from this study (supplementary table S3, Supplementary Material online). We retrieved the single copy-core proteins of the selected genomes per group for phylogenetic reconstruction: 351 protein group for *Buchnera* and 320 for *Erwinia*. We aligned the single-copy core protein sets using **MAFFT** v7.271 (Katoh and Standley, 2013) (L-INS-i algorithm). Divergent and ambiguously aligned blocks were removed using **Gblocks** v0.91b (Talavera and Castresana, 2007). The resulting alignments were concatenated for phylogenetic inference. Maximum-likelihood phylogenetic inferences were performed in **IQ-TREE** v1.6.8 using the LG+PMSF+R4 amino acid substitution model and ultrafast bootstrap approximation with 1000 replicates (Hoang *et al.*, 2018; Nguyen *et al.*, 2015; Wang *et al.*, 2018). LG was chosen since it incorporates the variability of evolutionary rates across sites in the matrix estimation (Le and Gascuel, 2008). The posterior mean site frequency (**PMSF**) model was chosen as it provides a rapid approximation to the profile mixture models C10 to C60 (variants of the CAT model available in PhyloBayes) (Wang *et al.*, 2018).

For phylogenetic placement of the novel *Hamiltonella* symbionts, we followed Chevignon *et al.* (2018) and used the *accD*, *dnaA*, *gyrB*, *hrpA*, *murE*, *ptsI*, and *recJ* gene sequences. We performed alignments using **MUSCLE** v3.8.31 (Edgar, 2004) and then removed divergent and ambiguously aligned blocks using **Gblocks** v0.91b. Bayesian inference was performed in **MrBayes** v3.2.5 (Ronquist *et al.*, 2012) using the GTR+I+G substitution model running two independent analyses with four chains each for 3,000,000 generations and checked for convergence.

To infer the origin of the Tn3 mobile elements found in *Erwinia* symbionts’ plasmids, we recovered similar sequences through on-line **BLASTP** searches *vs.* NCBI’s nr database and selected the top 50 non-redundant hits. Then, we aligned them using **MAFFT** v7.271 (L-INS-i algorithm) and removed divergent and ambiguously aligned blocks using **Gblocks** v0.91b. Bayesian inference was conducted in **MrBayes** v3.2.5 using the LG+G substitution model and ran as described above.

For phylogenetic inference of the putative horizontally-transferred genes, we collected orthologous genes across a selection of enterobacterial species (supplementary table S4, Supplementary Material online). Bayesian inference was performed in **MrBayes** v3.2.5 using the GTR+I+G substitution model and ran as previously described. Model selection for all nucleotide alignments was done in **jModelTest** v2.1.10 (Darriba *et al.*, 2012; Guindon and Gascuel, 2003).

All resulting trees were visualized and exported with **FigTree** v1.4.1 (http://tree.bio.ed.ac.uk/software/figtree/, last accessed February 7 2019) and edited in **Inkscape** v0.92.4. All alignments, NEXUS files, and NEWICK-/NEXUS-formatted trees can be found in https://doi.org/10.5281/zenodo.2566355 (last accessed February 7 2019).

## Results

### Localisation of *Cinara*-associated *Erwinia* symbionts in aphids

To investigate the localisation of the *Cinara*-associated *Erwinia* symbionts inside the aphid body, we performed FISH microscopy using competitive specific probes targeting the 16S rRNA from *Buchnera* and *Erwinia* symbionts of 3 species of aphids: *Cinara cuneomaculata* (fig. 1*A*), *Cinara kochiana* (fig. 1*B*), and *Cinara curvipes* (fig. 1*C*). Similarly to *Buchnera*, the *Erwinia* symbionts were found exclusively distributed inside bacteriocytes distinct from, but in close proximity to, those of *Buchnera*. They present a small coccoid shape contrasting with the larger round cells observed for *Buchnera*. This localisation resembles that of the *Se. symbiotica* symbionts from *C. cedri* and *Tuberolachnus salignus* (both holding a highly reduced genome) and contrasts that of the *Se. symbiotica* symbiont of *Cinara tujafilina* (holding a mildly reduced genome) (Manzano-Marín *et al.*, 2017).

**Figure 1.**
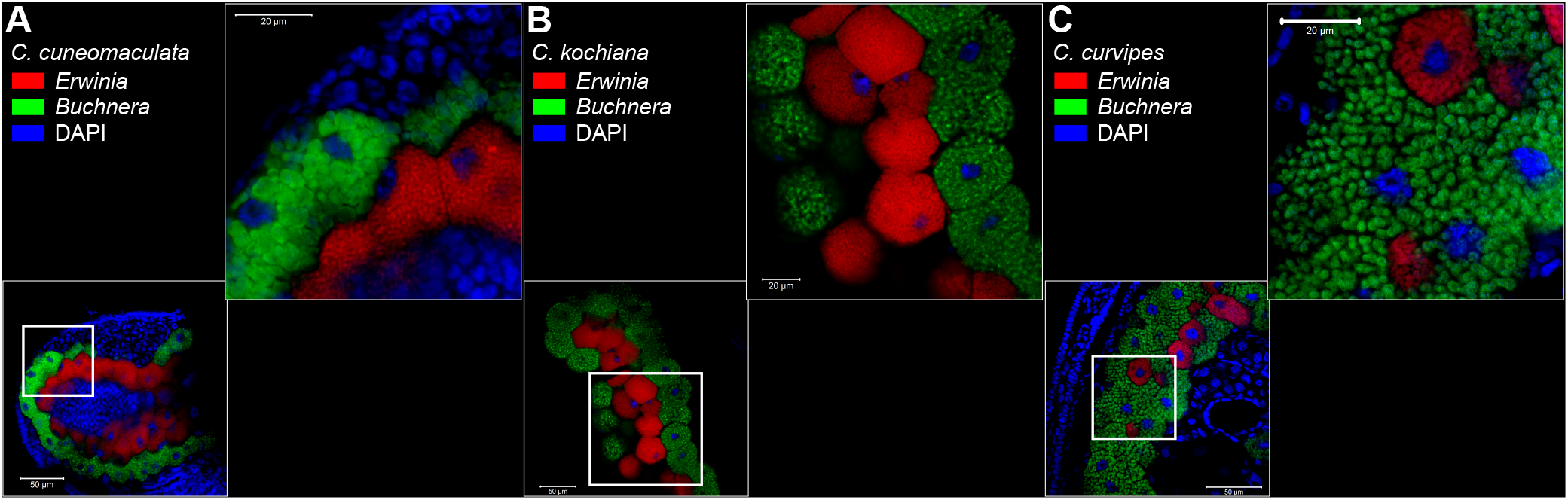
Location and morphology of *Erwinia* and *Buchnera* symbionts in selected *Cinara* aphids. Merged FISH microscopic images of aphid embryos from selected *Cinara* aphids. *Erwinia* signal is shown in red, *Buchnera*’s in green, and DAPI’s (staining DNA, highlighting host nuclei) in blue. Thick white boxes indicate the magnified region depicted in the top-right of each panel. The scientific name for each species along with the false colour code for each fluorescent probe and its target group are shown at the top-left of each panel. Unmerged images can be found in supplementary fig. S1 (Supplementary Material online).

### Phylogenetic history of the multi-partner endosymbiosis and *Erwinia* genome evolution

We successfully assembled the genomes of 9 *Buchnera*-*Erwinia* endosymbiont pairs from *Cinara* aphids. The *Buchnera* assemblies resulted in a single circular chromosome and two plasmids that code for leucine (pLeu, circularised) and tryptophan (pTrp, non-circularised) biosynthetic genes, respectively. The genomes of *Buchnera* are highly conserved in terms of number of genes, and have a size of between 442.57 and 458.25 kilo base pairs with an average G+C content of 23.01%. Average coverages range from 179-6410x (chromosome), 57-1252x (pLeu), and 843-9414x (pTrp) (supplementary table S5, Supplementary Material online). These genomes code for an average of 377 proteins, which are largely a subset of those coded by the previously sequenced *Buchnera* strains harboured by *C. cedri*, *C. tujafilina*, and *Cinara strobi*. Regarding the *Erwinia* endosymbionts, we recovered a circular chromosome and a circular plasmid for all strains, with average coverages ranging from 38-402x and 63-1682x, respectively.

A phylogenetic reconstruction using a set of single-copy core concatenated protein sequences shared across selected *Erwinia*, representing the currently-available diversity of the genus, and *Pantoea* species (fig. 2*A*). All *Cinara*-associated *Erwinia* endosymbionts (hereafter referred to as simply *Erwinia* endosymbionts) form a well-supported monophyletic group sister to that of a group of free-living *Erwinia* species that have been isolated as both pathogenic and non-pathogenic Rosaceae plant symbionts. When comparing the *Erwinia* endosymbionts’ tree topology *versus* that of *Buchnera* from the same aphid species, we observed a congruent evolutionary history between the two endosymbionts.

**Figure 2.**
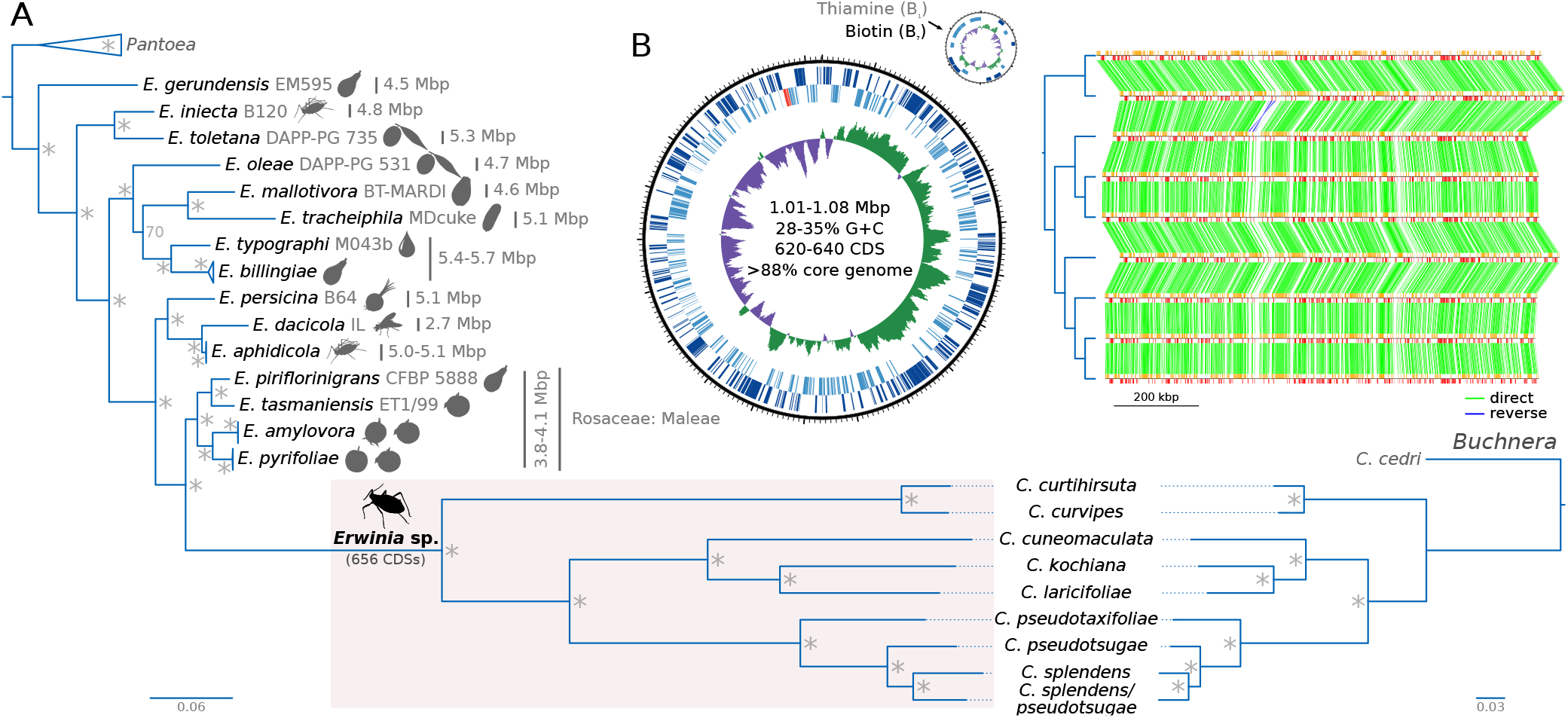
Phylogenetic reconstruction of *Erwinia* spp. and genome properties of *Cinara*-associated *Erwinia* endosymbionts. **(A)** On the left, phylogenetic reconstruction for *Erwinia* spp. using *Pantoea* spp. as an outgroup. *Erwinia* endosymbionts of *Cinara* species form a well supported monophyletic group. Silhouettes next to the leafs represents the isolation source for the strains. For the *Erwinia* symbionts, the name of the *Cinara* host is used. On the bottom right, the phylogeny of the corresponding *Buchnera* strains using the *Buchnera* associated with *C. cedri* as an outgroup is shown. Asterisks at nodes of trees stand for a posterior probability equal to 1. **(B)** On the left, the genome plot for the endosymbiont of *C. pseudotaxifoliae* along with statistics for sequenced *Erwinia* genomes. From outermost to innermost ring, the features on the direct strand, the reverse strand, and GC-skew plot are shown. On the right, pairwise synteny plots of *Erwinia* endosymbionts with a dendogram on the left displaying their phylogenetic relationships.

We found that the chromosomes of *Erwinia* endosymbionts all have a very similar genome size of around 1 Mbp, an increased A+T content of around 31.50% (when compared to free-living *Erwinia*), an average of 627 CDSs (including the plasmid-encoded proteins) (fig. 2*B*; supplementary table S6, Supplementary Material online), and no mobile elements. Regarding ncRNAs, they all code for a set of 36 tRNAs with charging potential for all of the 20 standard amino acids. A second copy of the *trnX* gene (tRNA-formylmethionine) is found in the symbionts of *C. kochiana* and *C. laricifoliae*, likely arising from a recent duplication event in the ancestor of these two species. All have one tmRNA gene, one 4.5S sRNA component of the Signal Recognition Particle, and the RNase P M1 RNA component. Each genome has an average of 14 pseudogenes, which are largely found as intact CDSs in other *Erwinia* endosymbiont strains. This, in combination with an analysis of shared genes among the *Erwinia* endosymbionts, revealed that the common ancestor of these strains coded for at least 656 distinct CDSs. We also identified a further event of gene duplication in the chromosome involving the *hflC* and *dapB* genes, occurring in the symbiont of *Cinara pseudotaxifoliae*. However, one copy of each gene has been pseudogenised, thus removing redundancy. Regarding genome organisation, the genomes of all sequenced *Erwinia* endosymbionts are highly syntenic (fig. 2*B*), with one major inversion of the region delimited by the *dapD* and *fldA* genes.

All *Erwinia* symbionts were found to have a putative multi-copy plasmid, which was recovered in the *Erwinia* bin during the assembly and binning process, regardless of the presence of additional symbionts. This plasmid mainly encodes for proteins involved in the biosynthesis of biotin (*bioA* and *bioB*) and thiamin (*thiC*, *thiF*, *thiS*, *thiG*, *thiH*, *thiD*, and *thiE*) as well as a FAD:protein FMN transferase (*apbE*), a 2,3-bisphosphoglycerate-dependent phosphoglycerate mutase (*gpmA*), a nucleoside permease (*nupC*), a putative heat shock protein (similar to IbpA/IbpB from *Escherichia coli*), and a PLD-like domain-containing protein. The thiamin-biosynthetic genes are notably missing in the plasmids of the *Eriwnia* symbionts of *Cinara curtihirsuta* and *C. curvipes*. It is important to note that most free living *Erwinia* retain an alternative thiamin-biosynthetic pathway, were they rely on the *thiO* gene (hosted in a plasmid along with the *thiSGF* genes) for the synthesis of iminoglycine. The exceptions to this rule are the genomes of *E. tasmaniensis* strain Et1/99 (*thiOSGF* in the chromosome) and the free-living *Erwinia* clade which includes *E. oleae*,where similarly to *Erwinia* endosymbionts, retain the *thiH* gene. The difference between the enzymes produced by these two genes is that the *thiH* gene uses L-tyrosine as precursor for iminoglycine, whereas *thiO* uses glycine.

Regarding their metabolic potential, *Erwinia* endosymbionts have a highly conserved metabolism (supplementary fig. S2, Supplementary Material online). They retain complete pathways for the synthesis of purine nucleotides from PRPP, L-glycine, and L-aspartate; while they retain salvage pathways for pyrimidines from thymidine and uridine. They encode for an intact FoF1-ATP synthase, a NADH:quinone oxidoreductase 1, a cytochrome *bo* oxidase, and a succinate:quinone oxidoreductase. They retain a fructose PTS permase and can perform glycolysis from glucose-1-P. In terms of the biosynthesis of the host’s essential amino acids (**EAAs**), and unlike their free-living relatives and *Buchnera*, *Erwinia* endosymbionts do not retain the capability to *de novo* synthesise for any of them, but keep specific importers for lysine, arginine, and threonine. The most intact route is that of L-lysine, where all strains lack the *lysA* gene and show differential retention of the *dapF* gene. Regarding vitamins and cofactors, *Erwinia* endosymbionts can synthesise riboflavin from GTP and ribulose-5P, biotin from KAPA, and thiamin pyrophosphate (the biologically active form of thiamin) from PRPP, L-cysteine, and L-tyrosine. This last vitamin biosynthetic pathway is notably missing from the endosymbionts of *C. curtihirsuta* and *C. curvipes*, which form a monophyletic clade. *Erwinia* endosymbionts retain the capability to synthesise and export peptidoglycan and (KDO_2_-lipid A). They preserve an intact Sec, Lpt, and Lol export systems as well as the porins OmpA and OmpC. Finally, they can *de novo* synthesise purines, but depend on the import or uridine, cytidine, and thymidine for the synthesis of pyrimidines.

### Metabolic complementation of bacterial partners and the tertiary co-obligate *Hamiltonella* symbiont

In previously analysed co-obligate nutritional endosymbiotic systems in Lachninae aphids, metabolic complementation is observed for some essential nutrients, including pathway splits regarding the biosynthesis of tryptophan and biotin (Manzano-Marín *et al.*, 2016) as well as the takeover of the riboflavin-biosynthetic role by *Se. symbitica* (Meseguer *et al.*, 2017). To infer the metabolic complementation between the *Buchnera* and *Erwinia* partners, we searched for the genes involved in the biosynthesis of EAAs, vitamins, and co-factors (fig. 3; supplementary figs. S3 and S4, Supplementary Material online). Similarly to *Se. symbiotica* co-obligate symbionts of Lachninae, the *Erwinia* endosymbionts retain an intact pathway for the biosynthesis of riboflavin as well as the *bioA*, *bioD*, and *bioB* genes; thus being able to complement *Buchnera*’s auxotrophy for these compounds. On the other hand, and as stated previously, they have lost most other EAAs’ biosynthetic genes. While most retain intact pathways for ubiquinol-8 and thiamin biosynthesis; the symbionts of *C. curtihirsuta* and *C. curvipes* have lost the *ubiI* gene as well as most thiamin-biosynthetic genes.

**Figure 3.**
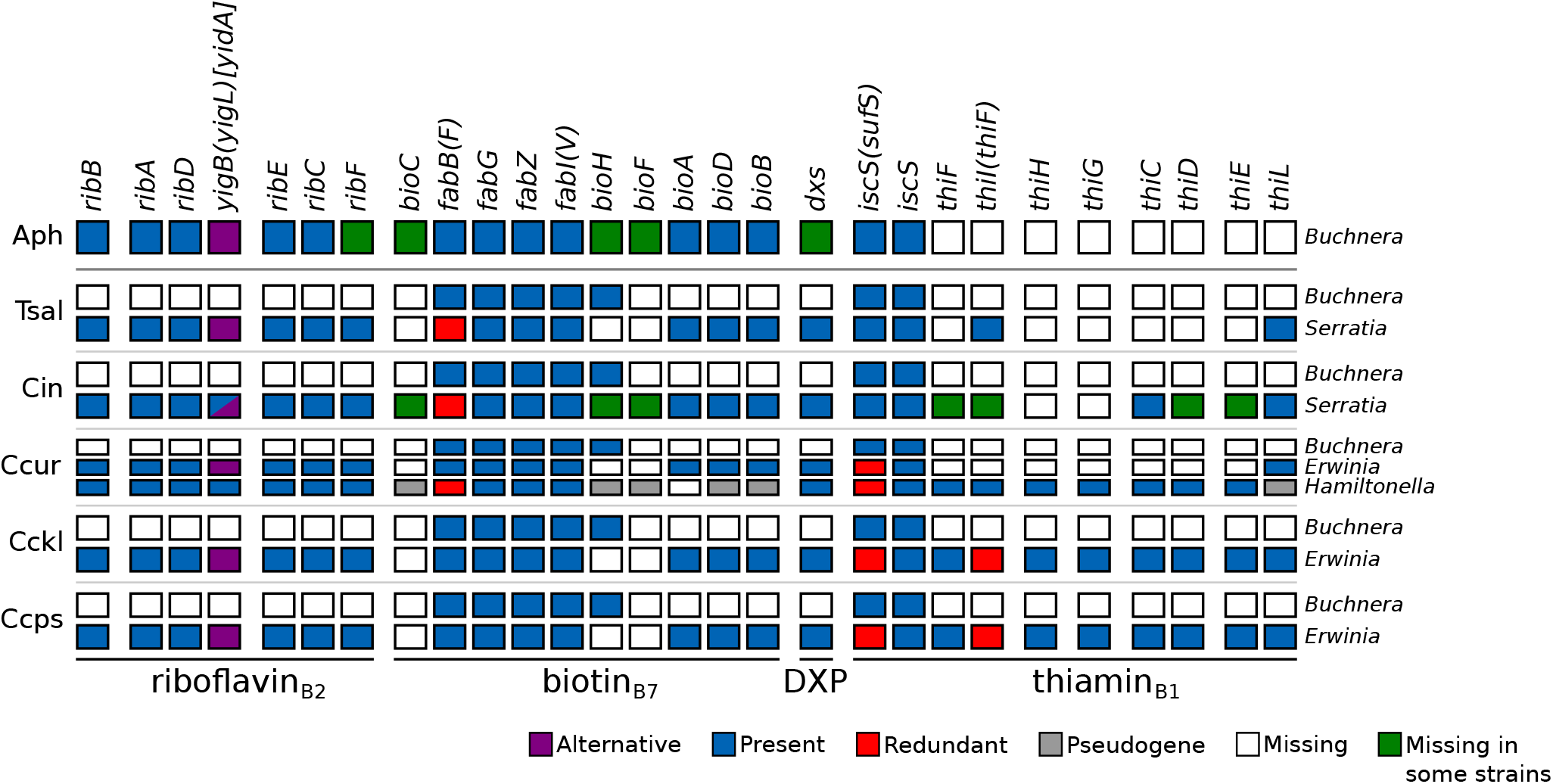
B-vitamin and cofactor biosynthetic metabolic complementation of obligate symbiotic consortia of different aphid species. Diagram summarising the metabolic complementarity of the fixed endosymbiotic consortia of co-obligate symbiotic systems of analysed *Erwinia*-associated *Cinara* aphids. For comparison, a collapsed representation of Aphididae *Buchnera*-only and Lachninae *Buchnera*-*Se. symbiotica* systems are used as outgroups. The names of genes coding for enzymes involved in the biosynthetic pathway are used as column names. Each row’s boxes represent the genes coded by a symbiont’s genome. At the right of each row, the genus for the corresponding symbiont. Abbreviations for the collapsed group of aphids harbouring the symbionts is shown at the left of each group of rows and goes as follows. Aph= Aphididae, Tsal= *T. salignus*, Cct= *C. cedri* + *C. tujafilina*, Ccur= *C. curtihirsuta* + *C. curvipes*, Cckl= *C. cuneomaculata* + *C. kochiana* + *C. laricifoliae*, Cpps= *C. pseudotaxifoliae* + *C. pseudotsugae* + *C. splendens* + *C.* cf. *splendens/pseudotsugae*. On the bottom, lines underlining the genes involved in the pathway leading to the compound specified by the name underneath the line.

Additionally, we have found a third bacterial partner from the *Hamiltonella* genus in both aforementioned *Cinara* species, which was previously determined to be a fixed symbiont in at least *C. curtihirsuta* and *C. curvipes* (Meseguer *et al.*, 2017). These *Hamiltonella* strains show highly syntenic reduced genomes of around 1.4 Mbp (supplementary fig. S5, Supplementary Material online), contrasting the 2.2 Mbp genomes of the *Hamiltonella defensa* facultative symbionts of *Ac. pisum* (Chevignon *et al.*, 2018). Based on 16S, their closest relatives are *H. defensa* strains ZA17 and AS3, with a 98.7% identity and a 2734 bit-score. Using the web server available at http://enve-omics.ce.gatech.edu/ani/ (last accessed May 22 2019) (Rodriguez-R and Konstantinidis, 2016), we calculated Average Nucleotide Identity (ANI) scores (Klappenbach *et al.*, 2007) for both *Cinara*-associated *Hamiltonella* symbionts *vs.* the facultative *H. defensa* symbiont strain ZA17, resulting in a two-way ANI score of 92.28% (SD: 3.53%) for the symbiont of *C. curtihirsuta* and 91.75% (SD: 3.79%) for that of *C. curvipes*, falling under the recommended species threshold (Chun *et al.*, 2018). However, a multi-gene phylogenetic reconstruction shows that these *Hamiltonella* symbionts are nested within the *H. defensa* symbiont clade (supplementary fig. S6, Supplementary Material online), confidently assigning these newly sequenced symbionts within the taxon. Based on a preliminary genome annotation of the assembled scaffolds, both *Hamiltonella* symbionts have around 940 CDSs, 50 possible pseudogenes, 38 tRNAs, 1 tmRNA, two copies of the rRNA operon, and 6 other ncRNAs (supplementary table S7, Supplementary Material online). The *Hamiltonella* symbionts associated with *Cinara* and *Erwinia* show a drastically reduced genetic repertoire, namely of CDSs and ncRNAs, but preserve a very similar G+C content to the facultative *H. defensa* (39% *vs.* 40%). Based on the single-copy core orthologous proteins of selected *H. defensa* symbionts (*Cinara*-associated and facultative strains ZA17, AS3, and 5AT), we observed very little rearrangement accompanied by a general loss of the plasmid and phage islands identified by Chevignon *et al.* (2018) (supplementary fig. S5, Supplementary Material online). Also, they lack the typical APSE phage found in facultative *H. defensa* strains. Regarding EAAs, vitamins, and co-factors, we performed an annotation and curation of the genes related to these pathways in the newly sequenced *H. defensa* and facultative *H. defensa* symbionts strains ZA17, AS3, and 5AT (supplementary fig. S7, Supplementary Material online). *Cinara*-associated *H. defensa* have the ability to code for L-threonine, lipoic acid, pyridoxal 5’-P (B_6_), riboflavin (B_2_), and thiamin (B_1_). Compared to the facultative *H. defensa* symbionts, they have lost the ability to synthesise biotin, having lost the *bioA* gene and undergoing pseudogenisation of the *bioC*, *bioH*, *bioF*, *bioD*, and *bioB* genes. They have also lost the capacity to synthesise chorismate (due to pseudogenisation and loss of all genes in the pathway) as well as ubiquinol-8, and L-lysine (due to the pseudogenisation of the *ubiC* and *lysA* genes).

### Serial horizontal gene transfer underlies multi-parter co-obligate mutualistic association

During the manual curation of the *Erwinia* endosymbiont plasmids, we noted that both the biotin- and thiamin-biosynthetic genes (*bioA*, *bioB*, *thiC*, *thiE*, *thiF*, *thiS*, *thiG*, *thiH*, and *thiD*) encoded in this molecule consistently showed top **BLASTP** hits against *Sodalis* and *Sodalis*-like bacteria, hinting at HGT. This contrasted with what we observed for the plasmid-encoded *nupC*, *apbE*, and *gpmA*, where the top **BLASTP** hits were consistently against *Erwinia* bacteria. The thiamin-biosynthetic genes present in the *Cinara*-associated *Hamiltonella*’s symbiont genomes also showed best BLAST hits against *Sodalis*-related bacteria. To test for HGT events across the *Erwinia* genome, we ran **BLASTP** similarity searches of the proteins of *Erwinia endosymbionts vs.* a database built from the proteomes of *Erwinia* and *Sodalis* species. The search and subsequent manual analysis revealed 11 proteins being from putative HGT origin from *Sodalis*-related bacteria: the plasmidic *bioA*, *bioB*, *thiC*, *thiE*, *thiF*, *thiS*, *thiG*, *thiH*, and *thiD* genes, and the chromosomal *bioD* and *thiI* genes. None of these genes were found to have a “native” copy in the genome that hosts them. To further test these putative HGT events, we collected orthologous genes across different enterobacterial species and reconstructed Bayesian phylogenies (fig. 4 and supplementary fig. S8, Supplementary Material online). All 11 genes showed strong support for a single event of HGT for both *Erwinia* and *Hamiltonella* symbionts of *Cinara*, consistently being recovered as a monophyletic group nested within or sister to *Sodalis* spp. This is in clear contrasts with what is observed for the putative non-HGT *gpmA* and *nupC* genes, that are confidently recovered nested within *Erwinia* spp. (supplementary fig. S8, Supplementary Material online). Additionally, the majority of the genes’ subtrees are congruent with the topology of the *Erwinia* endosymbionts’ subtree (supplementary fig. S9, Supplementary Material online). No *Sodalis*-related bacteria was detected during the assembly and binning process in any of the analysed samples.

**Figure 4.**
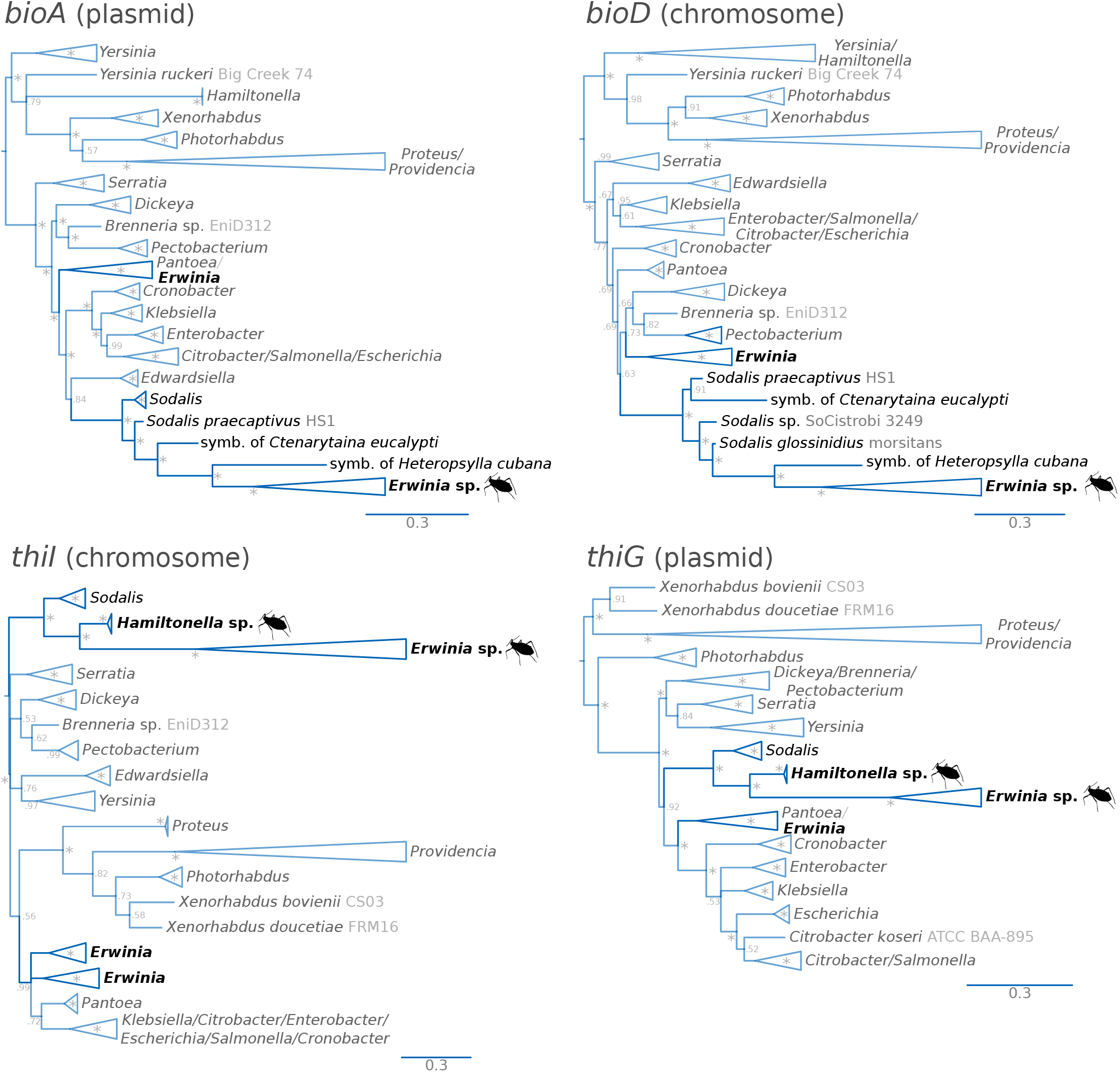
Phylograms of putatively horizontally transferred genes in *Erwinia* and *Hamiltonella* symbionts. Bayesian phylograms of four examples of the identified putative horizontally transferred genes present in the symbionts of the newly sequenced *Cinara* species. Taxon labels show the strain name in grey. Asterisks at nodes stand for a posterior probability equal to 1. On the bottom right, cladogram showing the relationships of genes congruent with the *Erwinia* phylogenetic tree.

To further explore the evolutionary history of the horizontally transferred genes, we analysed their location and gene order in the plasmids of *Erwinia* endosymbionts and the genomes of *Hamiltonella* symbionts and the free-living *Sodalis praecaptivus* (fig. 5). We chose this *Sodalis* strain since it is the only currently sequenced member of its genus that retains all thiamin-biosynthetic genes. Regarding the *Erwinia* endosymbionts’ plasmids, while they are very well conserved in terms of gene content, they display important changes in genome architecture. We observed that the symbionts of *C. cuneomaculata* and those of *C. pseudotaxifoliae*, *Cinara pseudotsugae*, *Cinara splendens*, and *Cinara* cf. *splendens/pseudotsugae* show the same gene order. However, within the monophyletic clade made up of the symbionts of *C. kochiana* and *Cinara laricifoliae*, we observed an ancestral duplication of the full gene repertoire. This has generated a plasmid with one intact and one pseudogenised copy of each gene in *C. kochiana* (except for the heat shock protein) and a new gene order in *C. laricifoliae*. All *Erwinia* endosymbionts belonging to the larger clade comprised of the aforementioned *Cinara* species show both *thiI* and *bioD* genes inserted into the same location in the chromosome. Within the monophyletic cluster made of *C. curtihirsuta* and *C. curvipes*, the *Erwinia* endosymbionts show a much different plasmid architecture than those of their sister clade. Their plasmid shows an inverted duplication which includes the *bioA* and *bioB* genes as well as the genes coding for the PLD-like domain-containing protein, a putative heat shock protein, a replication-associated protein (*repA2*), and the Tn3 family resolvase/invertase (a mobile element). The presence of the latter is puzzling, given that the other sequenced *Erwinia* endosymbionts completely lack mobile elements. A Bayesian phylogenetic analyses revealed that this Tn3 family resolvase/invertase is most closely related to Tn3 resolvase/invertases encoded by other *H. defensa* symbionts (supplementary fig. S10, Supplementary Material online), hinting at their origin. As mentioned before, the *Erwinia* endosymbionts of *C. curtihirsuta* and *C. curvipes* do not host the “typical” thiamin-biosynthetic genes in their plasmid. In turn, these genes are now hosted in the *Hamiltonella* genome, flanked by insertion sequence (hereafter **IS**) elements and a glycerol dehydrogenase pseudogenes. Whether or not these genes reside in a plasmid remains unclear. However, the scaffolds in which these genes reside in each *Cinara*-associated *Hamiltonella* strain, and which bear no resemblance to *Erwinia* plasmids, has roughly triple the coverage as the rest of the large scaffolds (*circa* 1,900 *vs.* 5,400x and 620x *vs.* 1,500x), suggesting either a triplication in the chromosome or their presence in a multi-copy plasmid of at least 42.8 kbp in the symbiont of *C. curtihirsuta* and 73.2 kbp in the one of *C. curvipes*. The plasmidic nature of these molecules is supported by the presence of a replication initiator protein gene and a plasmid segregation protein in the assembled sequences for both *Hamiltonella* strains. When comparing the genomic context of the horizontally transferred genes against that of the same genes in *So. praecaptivus*, several rearrangements have taken place. Namely, the *thiD* gene is now hosted by the aphid symbionts in close proximity to the rest of the “*thi*” genes and the *bioD* gene has been distanced from the other biotin-biosynthetic genes. In all cases, the *thiL* gene, coding for the last enzymatic step in the biosynthesis of thiamin diphosphate (the active form *in vivo*), is located solely in the chromosome of *Erwinia* endosymbionts.

**Figure 5.**
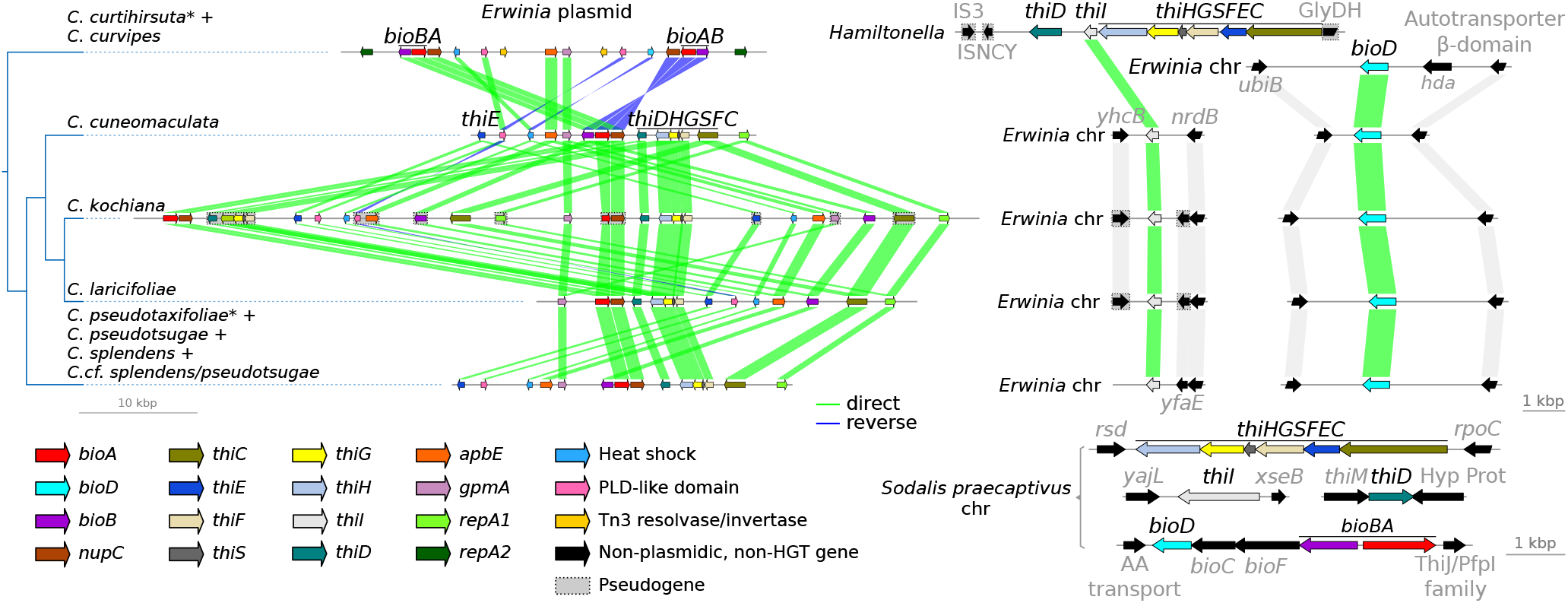
Evolution of the horizontally-transferred biotin- and thiamin-biosynthetic genes. Linear plots of the genomic context for the genes originating from a HGT from *Sodalis*-related bacteria. On the left, dendogram showing the phylogenetic relations of the different *Erwinia*-associated *Cinara* groups of species. On the bottom-right, genomic context for the aforementioned genes in *So. praecaptivus* for comparison. chr= chromosome.

### ’*Candidatus* Erwinia haradaeae’ sp. nov

We propose the specific name ’*Candidatus* Erwinia haradaeae’ for the monophyletic lineage of enterobacterial endosymbionts exclusively found affiliated as co-obligate symbionts in the monophyletic group of *Cinara* aphid species analysed in this study, although its presence in other closely related species cannot be discarded. The closest relative of the symbiont of *C. pseudotaxifoliae* by 16S rRNA gene sequence identity is the *Erwinia pyrifoliae* strain EpK1/15 (INSDC accession number CP023567.1) with which it shares 94 % sequence identity (2350 bit-score). The specific epithet ’haradaeae’ is in honour of Hosami Harada, who performed research suggesting the origin of the aphid’s obligate *Buchnera* symbiont as derived from a habitant of the plant on which the host fed (Harada and Ishikawa, 1993; Harada *et al.*, 1996). These studies put forward the idea of the aphids’ symbionts possibly evolving from bacteria originally inhabiting the plant, then transitioning to gut associates, and finally obligate symbionts.

In *C. cuneomaculata*, *C. kochiana*, and *C. curvipes*, ’*Ca.* Erwinia haradaeae’ is found inhabiting the bacteriome inside bacteriocytes distinct from those of *Buchnera*. Previous work referring to these symbionts as ’*Erwinia*’ or ’*Erwinia*-related’ include Jousselin *et al.* (2016) and Meseguer *et al.* (2017). Currently available sequences that correspond to ’*Ca.* Erwinia haradaeae’ are deposited under INSDC accession numbers LT670851.1 and LT670852.1. All currently known ’*Ca.* Erwinia haradaeae’ species harbour biotin- and thiamin-biosynthetic horizontally transferred genes from a *Sodalis* or *Sodalis*-related bacteria both in their chromosome and plasmid. The exception to this rule are the strains associated with a co-obligate *Ca.* Hamiltonella defensa symbiont, where the thiamin-biosynthetic genes of horizontal-transfer origin are missing, and in turn are located in a putative plasmid of *Ca.* Hamiltonella defensa. Based on genome-based metabolic inference and their parallel evolutionary history with *Buchnera*, they all are co-obligate endosymbionts along with *Buchnera*, and *Ca.* Hamiltonella defensa in *C. curtihirsuta* and *C. curvipes*, in the *Cinara* species sequenced and presented in this study.

## Discussion

Obligate nutritional microbial endosymbioses are among the most integrated forms of beneficial associations between eukaryotes and bacteria. Obligate symbiotic systems that involve more than one co-obligate nutritional partner have not only been described in aphids, but also in other hemipterans such as planthoppers (Bennett and Mao, 2018), cicadas (McCutcheon *et al.*, 2009; Van Leuven *et al.*, 2014), scale insects (Husnik *et al.*, 2013; Szabó *et al.*, 2017), among others (McCutcheon and Moran, 2007; Weglarz *et al.*, 2018). These associations have proven to be quite dynamic in certain lineages, with several events of symbiont replacement occurring along their evolutionary history (Husnik and McCutcheon, 2016; Matsuura *et al.*, 2018; Toenshoff *et al.*, 2012; von Dohlen *et al.*, 2017). The *Cinara* aphids offer a great opportunity to study the mechanisms of symbiont turnover: they display a rather unstable multi-partner co-obligate symbiosis with frequent events of secondary symbiont replacement well-supported by a reliable host phylogeny. In this work we have analysed the recently evolved co-obligate association between *Buchnera* and *Ca.* Erwinia haradaeae (hereafter *E. haradaeae*) in order to assess its origin and evolutionary stabilit in an otherwise dynamic mutualistic partnership.

Through phylogenetic and genome rearrangement analyses of *Erwinia* endosymbionts’ genomes, we have determined that these symbionts most likely arose, similarly to *Buchnera*, from a single event of host fixation and have been co-diverging with *Buchnera* ever since. Given the age of the aphid clade hosting these symbionts (Meseguer *et al.*, 2015), these new *Erwinia* obligate mutualistic partners have co-diverged with their aphid hosts and associated *Buchnera* for about 20-25 million years. Like their co-obligate *Buchnera* partner, and mirroring the genomic evolution of distantly related co-obligate nutritional symbionts, *E. haradaeae* strains reside in bacteriocytes (close to *Buchnera*) and have evolved a highly reduced genome mostly deprived of mobile elements. This work provides strong support for an evolutionary history involving an early drastic genome reduction and loss of mobile elements and an intracellularisation of *E. haradaeae* in the ancestor of their *Cinara* hosts’. This chain of events mirrors the one inferred for other obligate endosymbionts (Pérez-Brocal *et al.*, 2006; Williams and Wernegreen, 2015). Regarding their evolutionary origin, we found that *E. haradaeae* is most closely related to a group of *Erwinia* species that have been isolated as both pathogenic and non-pathogenic plant symbionts. *Erwinia* bacteria are generally described as plant-associated bacteria (Kado, 2006). Nonetheless, *Erwinia* species have also been isolated from insects, namely the olive fruit fly (*Bactrocera oleae*) (Capuzzo, 2005), the western flower thrips (*Frankliniella occidentalis*) (de Vries *et al.*, 2001) and aphids (Fischer-Le Saux *et al.*, 2015; Harada and Ishikawa, 1993; Harada *et al.*, 1996, 1997). Concerning the latter, *Erwinia* was isolated both from artificial diets exposed to probing of *Diuraphis noxia* biotype 2 sylets through a Parafilm membrane (*Erwinia iniecta*) and from the gut of a lab strain of *Ac. pisum* (*Erwinia aphidicola*). The uptake and persistence of this bacteria within the aphid digestive tract is supported by an early experiment demonstrating that the aphid *Aphis pomi* was capable of ingesting a pathogenic *Erwinia amylovora* after a short feeding period (5 minutes) and this would persist in the bodies of the insects for at least 72 hours (duration of the longest experiment) (Plurad *et al.*, 1965). Thus, aphid feeding is a likely source for the presence and persistence of *Erwinia* bacteria inside the aphid digestive tract, and would facilitate a frequent and prevalent interaction between *Erwinia* strains and aphids. It is important to note that the aforementioned *Erwinia* symbionts described for *F. occidentalis* (Facey *et al.*, 2015) belong to the *E. aphidicola* species (AB681773.1; type strain X 001=CIP 106296=IAM 14479=JCM 21238=LMG 24877=NBRC 102417), based on 16S sequence identity (between 99.72-100.00%). This indicates that strains from this bacterial species can actually infect related insect species such as aphids and thrips (Thysanoptera: Thripidae).

As suggested in Meseguer *et al.* (2017), *E. haradaeae* have evolved a very compact genome that has lost the capacity to synthesise any essential amino acid (with respect to host’s requirements; supplementary fig. S2, Supplementary Material online), but have retained metabolic pathways for complementing *Buchnera* in regards to the biosynthesis of two B vitamins: biotin and riboflavin. This pattern is consistent with what is observed in two independently-evolved *Buchnera*-*Se. symbiotica* co-obligate systems found Lachninae aphids (Manzano-Marín *et al.*, 2016). It shows an effective partitioning of essential nutrient biosynthesis that has probably partly evolved through a three-way coevolutionary process between aphids and their two bacteria. Though the intricacies of metabolite exchanges in this system remain to be discovered, the separation of *Buchnera* and *E. haradaeae* into distinct bacteriocytes implies that most interactions between the two endosymbionts are mediated by the hosts. In addition, and unlike the previously analysed nutritional “di-symbiotic” systems from Lachninae and “mono-symbiotic” ones from Aphidinae aphids, we found that the capacity to synthesise thiamin, another essential B vitamin, was conserved in all newly sequenced symbiotic consortia. Regarding this vitamin, it was found that *Neomyzus circumflexus* (Aphidinae) aphids fed on an artificial diet missing thiamin, mortality in the first generation was normal at around 5% and the adults were found to be much smaller and gave 54% less larvae than the control animals (Ehrhardt, 1968). Additionally, the development of the 2nd generation was stagnating, being the larvae stuck in the 2nd instar after 20 days. Taken together, these results support the essential nature of this vitamin and suggest that the diet from the *E. haradaeae*-associated *Cinara* might have a lower concentration of thiamin than that of the previously sequenced aphid symbiotic systems. To our knowledge, no studies have looked at sap vitamin contents in the conifer genera on which this monophyletic group of aphids feed (i.e. mainly *Pseudotsuga*, *Larix*, *Abies*, and a few *Picea* species; see Jousselin *et al.* 2013 and Meseguer *et al.* 2015 for studies on aphid host plant association in this clade). In addition to analysing the phloem of these trees, it would be valuable to investigate the presence of thiamin-biosynthetic genes in aphid species feeding on the same host plants. The *E. haradaeae*-associated *Cinara* group actually contains the only aphid species feeding on *Larix*. Nonetheless, other *Cinara* species also feed on *Abies*, *Picea*, and *Pseudotsuga*. The endosymbiont and aphid host genomes of these species would be worth investigating.

Most strikingly, we found strong evidence supporting the horizontal transfer of not only three biotin-biosynthetic genes (*bioA*, *bioD*, and *bioB*), but also most thiamin-biosynthetic ones (except for *thiL*). Through phylogenetic analyses, we were able to confidently determine that the likely origin of the HGT genes in *Erwinia* endosymbionts and *Hamiltonella* was a *Sodalis* or *Sodalis*-related bacterium. *Sodalis*-allied bacteria are found across different insect taxa as facultative or obligate endosymbionts (Koga and Moran, 2014; Oakeson *et al.*, 2014; Santos-Garcia *et al.*, 2017; Sloan and Moran, 2012) and likely in plant tissues (Chari *et al.*, 2015). Furthermore, bacteria from these taxa have been previously identified to be associated with aphids (Manzano-Marín *et al.*, 2017, 2018; Meseguer *et al.*, 2017). Therefore, it is likely that the co-inhabitation of *Erwinia* and a *Sodalis*-allied bacterium facilitated the horizontal transfer of these genes to the plasmid of *E. haradaeae* with subsequent transfers of *bioD* and *thiI* to the chromosome. HGT involving genes in vitamin biosynthetic pathways has been previously documented in *Candidatus* Legionella polyplacis, the endosymbiont of the blood-feeding louse *Polyplax serrata* (Říhová *et al.*, 2017). This *Legionella* symbiont has acquired a complete biotin operon (*bioADCHFB*) likely from a rickettsial bacterium. A similar example is observed in the *Cardinium* endosymbionts of the parasitoid wasp *Encarsia pergandiella* (Penz *et al.*, 2012) and the whitefly *Bemisia tabaci* (Říhová *et al.*, 2017). The localisation of these genes to a plasmid is not surprising, with examples of other nutritional endosymbionts such as *Buchnera* or *Riesia*, where genes important for their symbiotic role as nutrient providers are hosted in such extrachromosomal elements (Boyd *et al.*, 2014; Bracho *et al.*, 1995; Gil *et al.*, 2006; Kirkness *et al.*, 2010; Lai *et al.*, 1994). Such phenomenon has been proposed to be an adaptation to overexpression in *Buchnera* (Lai *et al.*, 1994), and it would be consistent with our findings of both the *Erwinia* plasmids and the *Hamiltonella* contigs (putatively plasmidic) bearing the *bio* and/or *thi* genes to be amplified relative to the chromosome.

While most *E. haradaeae* retain the thiamin-biosynthetic genes, the monophyletic group of *Erwinia* endosymbionts from *C. curtihirsuta* and *C. curvipes* have lost all horizontally-transferred thiamin-biosynthetic genes. Our results suggest that these genes have been further horizontally-transferred to a tertiary co-obligate *H. defensa* symbiont, thus locking a new tri-symbiotic co-obligate nutritional symbiosis. These genes are located in a large putative multi-copy plasmid in both *Hamiltonella*, analogous to what is observed for *E. haradaeae*. The handing on of the thiamin-biosynthetic genes from *E. haradaeae* to yet a newcomer, *Hamiltonella*, that has now become obligate, further supports the hypothesis that these genes have a functional significance for the mutualistic association between aphids and their consortium of bacteria. The new *Hamiltonella* symbiont preserves genes to synthesise pyridoxal-5P (vitamin B_6_). The ability to synthesise this compound is consistently missing in all other mono- and di-symbiotic systems, and thus could represent either a remnant from the genome reduction process in the co-obligate *Hamiltonella* or an important addition to sustain either itself or the tri-symbiotic endosymbiotic consortium (due to the putative increased demand for this compound). Although Meseguer *et al.* (2017) suggested that the replacement of co-obligate symbionts in *Cinara* might not be adaptive for the aphid hosts (they do not correlate with broad ecological niche shifts of aphids), the fact these new symbionts carry and retain genes with new metabolic functions might suggest otherwise. The shift in ecological niche enabled by the replacement of symbionts alongside *Buchnera*, might be detectable at a finer scale than the one explored by Meseguer *et al.* (2017); it could involve fine tuned adaptations to phloem sap nutrient content.

Altogether, our results allow us to propose an evolutionary scenario for the *Buchnera*-*Erwinia* co-obligate symbiotic consortium in *Cinara* aphids (fig. 6). First, the *Erwinia* bacterium associated with aphids, likely originating from a plant-affiliated symbiont, replaced the previously existing secondary co-obligate symbiont of *Buchnera*. The former secondary co-obligate symbiont was likely a *Se. symbiotica* bacterium, as suggested by an ancestral reconstruction of the bacterial symbiotic associations within *Cinara* aphids (Meseguer *et al.*, 2017). The recruitment of a new obligate symbiont among bacteria likely found in the insect feeding source has not been commonly documented(Douglas, 2016), and represents a novel finding in evolutionary scenarios of multi-partner endosymbioses. The ability to replace the former co-obligate symbiont was facilitated by the presence of horizontally transferred biotin- and thiamin-biosynthetic genes in its plasmid from a co-existing *Sodalis* symbiont. Subsequently, the gene *bioD* was transferred into the chromosome of *E. haradaeae*. In addition to intra-cellularisation, *E. haradae* underwent a drastic genome reduction, loosing in the process most EAA and B vitamin biosynthesis genes, except for those needed to assist *Buchnera*. This was followed by 20 million years of genome stasis and codivergence with its partner. Afterwards, in the lineage leading to *C. curtihirsuta* and *C. curvipes*, *E. haradaeae* lost the capacity to synthesise thiamin due to the loss of the *thi* genes. Nonetheless, this loss was accompanied by the acquisition by HGT of the aforementioned genes from *E. haradaeae* to the co-existing *H. defensa*. This second HGT event locked the tri-symbiotic system together. In the meantime, in the lineage leading to the sister clade of *Hamiltonella*-associated *Cinara* aphids, *E. haradaeae*’s genome underwent a further event of plasmid to chromosome transfer of the gene of HGT origin *thiI*.

**Figure 6.**
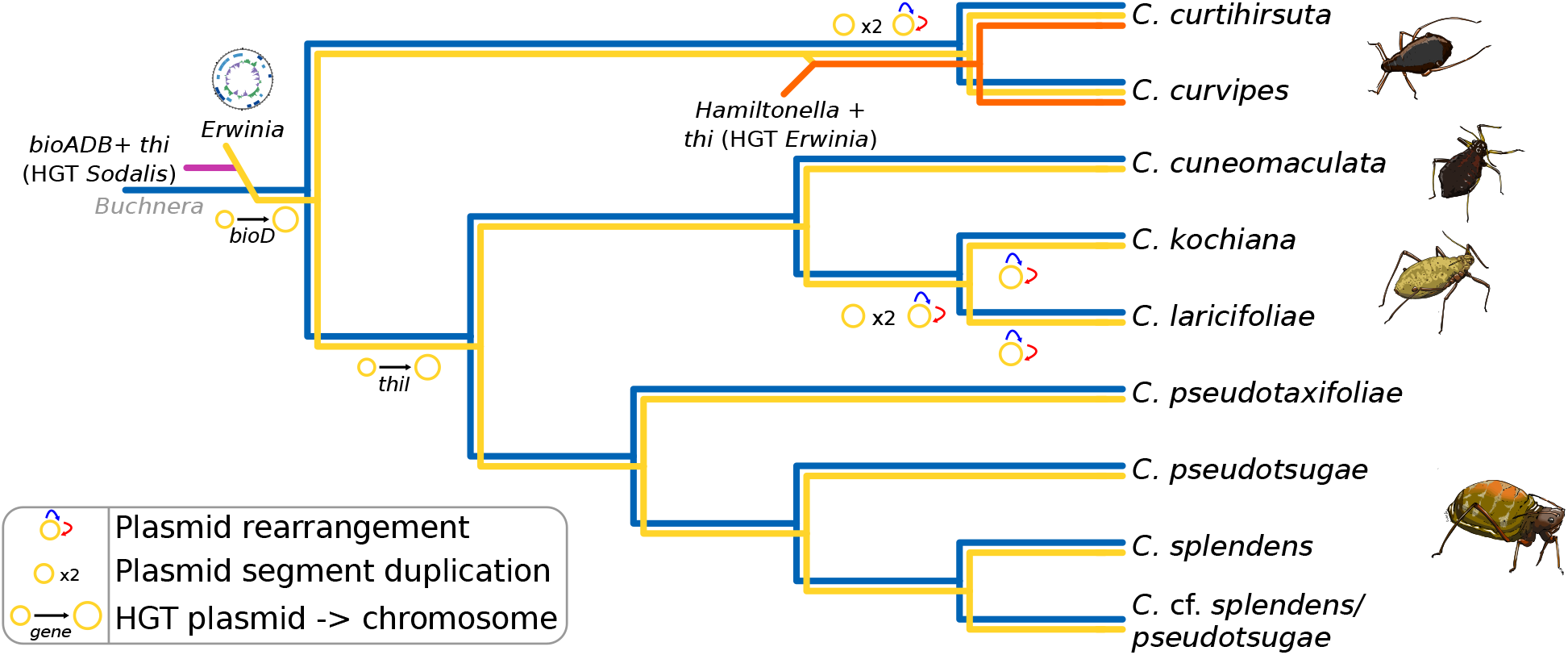
Proposed evolutionary scenario for the establishment and evolution of co-obligate symbionts of *E. haradaeae*-associated *Cinara*. Cladogram displaying the relationships of sequenced *E. haradaeae*-associated *Cinara* lineages. Coloured lines in the dendogram are used to represent the persistence of a symbiont in the aphid lineage. Incoming lines on branches symbolise the acquisition of co-obligate secondary symbionts. Lines joining two symbiotic lineages symbolise HGT events. At the leaves, cartoons of selected aphids form the different groups of genera are shown.

## Conclusion

Symbiont replacement is a phenomenon observed in multi-partner endosymbioses (Husnik and McCutcheon, 2016; Koga and Moran, 2014; Toenshoff *et al.*, 2012), being the newcomer the one that is often repeatedly replaced. The presented results reveal the crucial role that horizontal transfer of pivotal metabolic genes across co-existing symbionts can have in this dynamics. First explored by Harada *et al.* (1996), the current work also provides support for the bacteria present in the aphid’s diet as being a likely source of obligate symbiotic bacteria. Finally, this work also raises new questions about the role of these new symbionts in aphids’ ecological niches. While acquisitions and replacements of bacteria in mutli-partner endosymbioses do not generally involve the recruitment of new nutritional functions (Douglas, 2016), here our results suggest otherwise. Indeed, our genomic enquiry suggests that thiamin biosynthesis genes might be pivotal in the establishment of both *E. haradaeae* and then *Hamiltonella*, raising questions about the importance of this nutrient for the insect host and therefore the role of host level selection in the process of symbiont replacement. We expect that further exploration of the intricacies of the di-symbiotic *Buchnera*-*Erwinia* and tri-symbiotic *Buchnera*-*Erwinia*-*Hamiltonella* systems will continue to provide important clues into the emergence and maintenance of these multi-partner endosymbioses.

## Acknowledgements

We would like to acknowledge the talented artist/scientist Jorge Mariano Collantes Alegre for the aphid cartoons in fig. 6. This work was supported by the *Marie-Curie AgreenSkills+* fellowship programme co-funded by the *EU’s Seventh Framework Programme* (FP7-609398) to A.M.M., the *Agropolis foundation/Labex Agro* (”*Cinara*’s microbiome”) to E.J, the the *France Génomique* National Infrastructure, funded as part of the *Investissemnt d’Avenir* program managed by the *Agence Nationale pour la Recherche* (ANR-10-INBS-09) to C.O, C.C., and V.B. This publication has been written with the support of the AgreenSkills+ fellowship programme which has received funding from the EU’s Seventh Framework Programme under grant agreement No. FP7-609398 (AgreenSkills+ contract). We are grateful to the genotoul bioinformatics platform Toulouse Midi-Pyrenees (Bioinfo Genotoul) for providing help and/or computing and/or storage resources. The authors are grateful to the CBGP-HPC computational platform. The funders had no role in study design, data collection and analysis, decision to publish, or preparation of the manuscript.

