## Supplementary material for "Serial horizontal transfer of vitamin-biosynthetic genes enables the establishment of new nutritional symbionts in aphids’ di-symbiotic systems": File S2

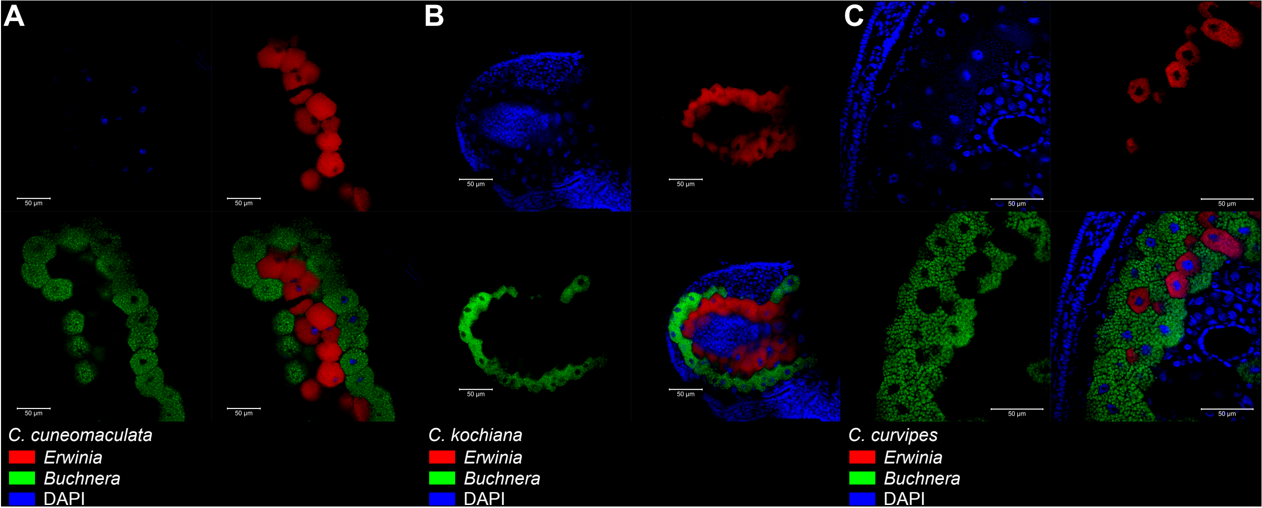


**Fig. S1. Location and morphology of *Erwinia* and *Buchnera* symbionts in selected *Cinara* aphids.** FISH microscopic images of aphid embryos from selected *Cinara* aphids. Symbiont-specific probes were used for FISH. *Erwinia* is shown in red, *Buchnera* in green, and the DAPI signal is shown in blue. The scientific name for each species along with the false colour code for each fluorescent probe and its target group are shown at the top-left of each panel. Scale bars from the unmagnified and magnified FISH images are shown as white bars.

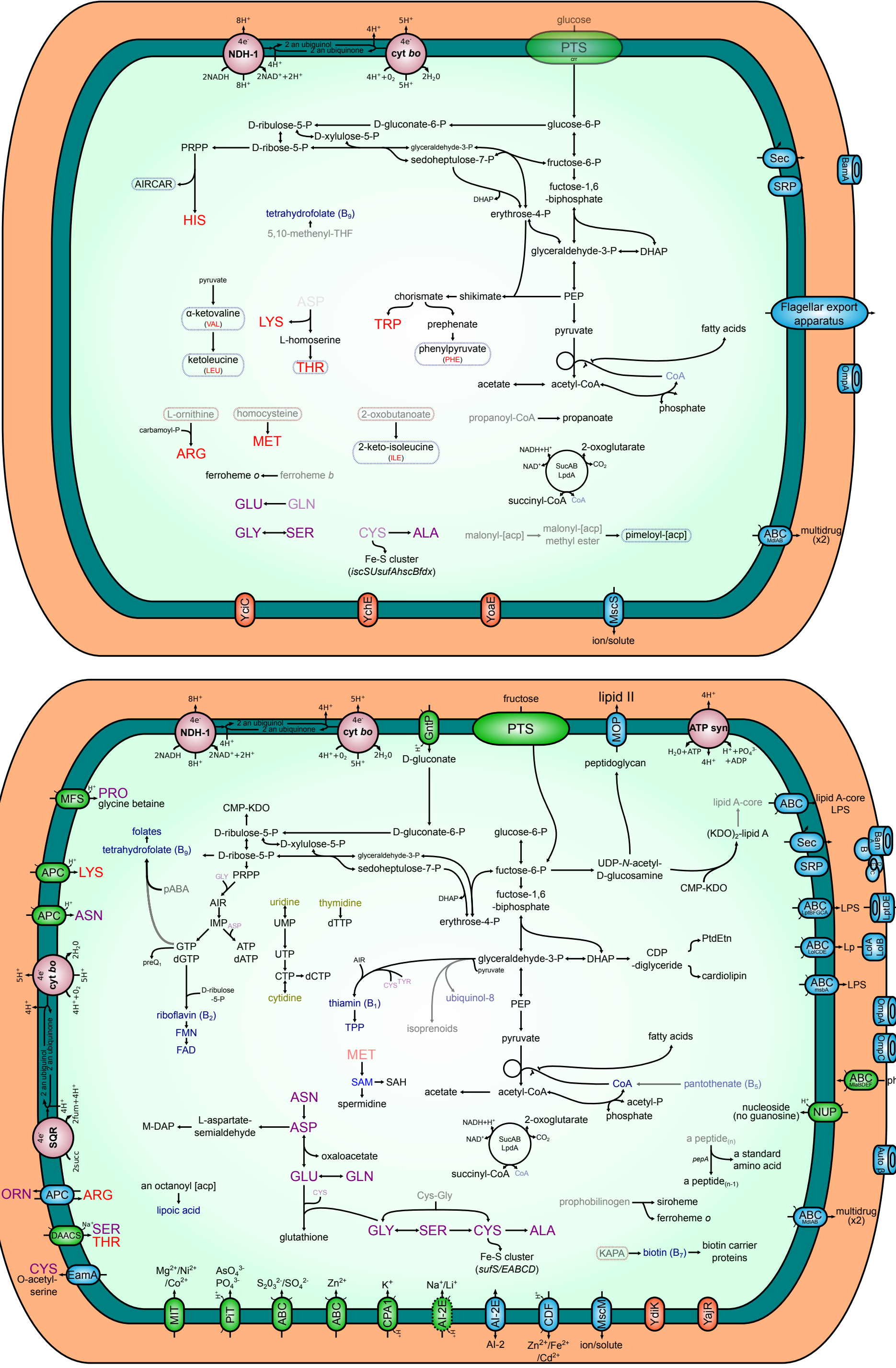


**Fig. S2. Metabolic reconstruction of *Buchnera-Erwinia* disymbiotic system of *Cinara pseudotaxifoliae*.** Intact pathways are represented with solid black lines, and unclear ones (missing a specific gene or having it pseudogenized by a frameshift) with solid gray lines. Importers are displayed using green ovals, while exporters and exporters/importers are displayed using blue ovals. The name inside each oval states the family/superfamily they belong to (following TCDB's classification [Saier *et al.* 2014]), otherwise the protein name is used. Essential and nonessential amino acids are shown in red and purple lettering, respectively, while cofactors and vitamins are shown in blue. Blurred compounds represent those for which biosynthesis or import cannot be accounted for based on genomic data. Blurred transporters represent those for which a part of the transporter is missing, therefore recently pseudogenized. In *Buchnera*, dotted boxes indicate identified imported/exported compounds for the biosynthesis of amino acids and nucleotides. The symbionts of *C. pseudotaxifoliae* were used as representatives from *Erwinia-Buchnera* endosymbiotic systems.

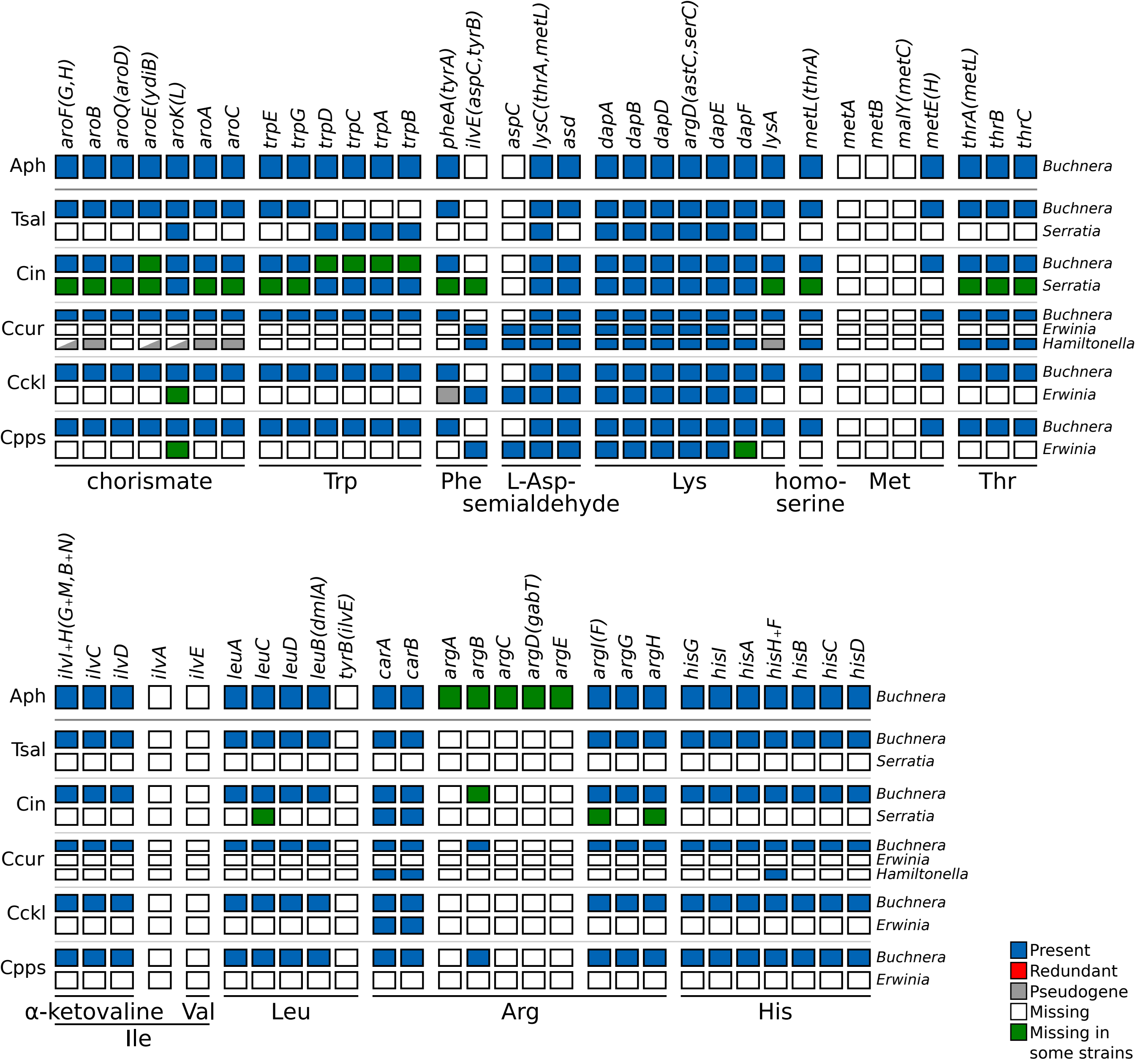


**Fig. S3. Essential amino acid biosynthetic metabolic capabilities of obligate symbiotic consortia of different aphid species.** Diagram summarising the metabolic capabilities of the fixed endosymbiotic consortia of co-obligate symbiotic systems of analysed *Erwinia*-associated *Cinara* aphids. For comparison, a collapsed representation of Aphididae *Buchnera*-only and Lachninae *Buchnera-Serratia symbiotica* systems are used as outgroups. The names of genes coding for enzymes involved in the biosynthetic pathway are used as column names. Each row's boxes represent the genes coded by a symbiont's genome. At the right of each row, the genus for the corresponding symbiont. Abbreviations for the collapsed group of aphids harbouring the symbionts is shown at the left of each group of rows and goes as follows. Aph= Aphididae, Tsal= *T. salignus*, Cct= *C. cedri* + *C. tujafilina*, Ccur= *C. curtihirsuta* + *C. curvipes*, Cckl= *C. cuneomaculata* + *C. kochiana* + *C. laricifoliae*, Cpps= *C. pseudotaxifoliae* + *C. pseudotsugae* + *C. splendens* + *C. cf. splendens/pseudotsugae*. On the bottom, lines underlining the genes involved in the pathway leading to the compound specified by the name underneath the line.

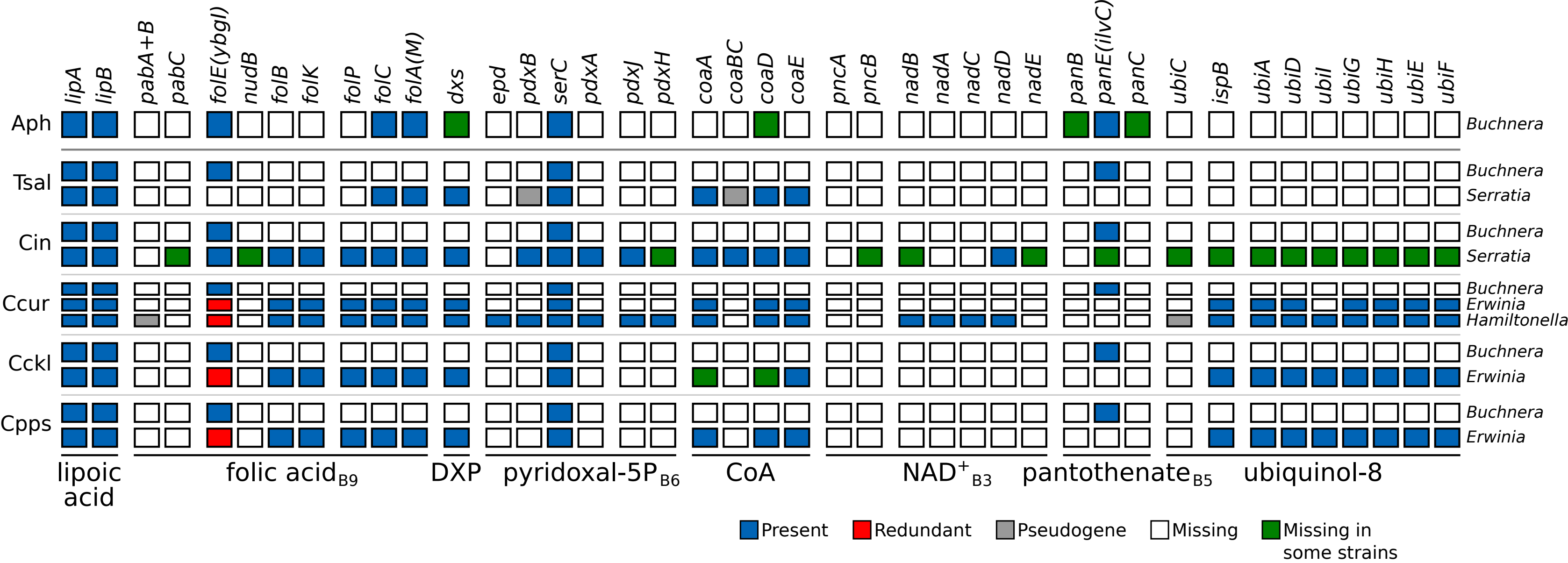


**Fig. S4. Supplementary B-vitamin and cofactor biosynthetic metabolic capabilities of obligate symbiotic consortia of different aphid species.** Diagram summarising the metabolic capabilities of the fixed endosymbiotic consortia of co-obligate symbiotic systems of analysed *Erwinia*-associated *Cinara* aphids. For comparison, a collapsed representation of Aphididae *Buchnera*-only and Lachninae *Buchnera-Serratia symbiotica* systems are used as outgroups. The names of genes coding for enzymes involved in the biosynthetic pathway are used as column names. Each row's boxes represent the genes coded by a symbiont's genome. At the right of each row, the genus for the corresponding symbiont. Abbreviations for the collapsed group of aphids harbouring the symbionts is shown at the left of each group of rows and goes as follows. Aph= Aphididae, Tsal= *T. salignus*, Cct= *C. cedri* + *C. tujafilina*, Ccur= *C. curtihirsuta* + *C. curvipes*, Cckl= *C. cuneomaculata* + *C. kochiana* + *C. laricifoliae*, Cpps= *C. pseudotaxifoliae* + *C. pseudotsugae* + *C. splendens* + *C. cf. splendens/pseudotsugae*. On the bottom, lines underlining the genes involved in the pathway leading to the compound specified by the name underneath the line.

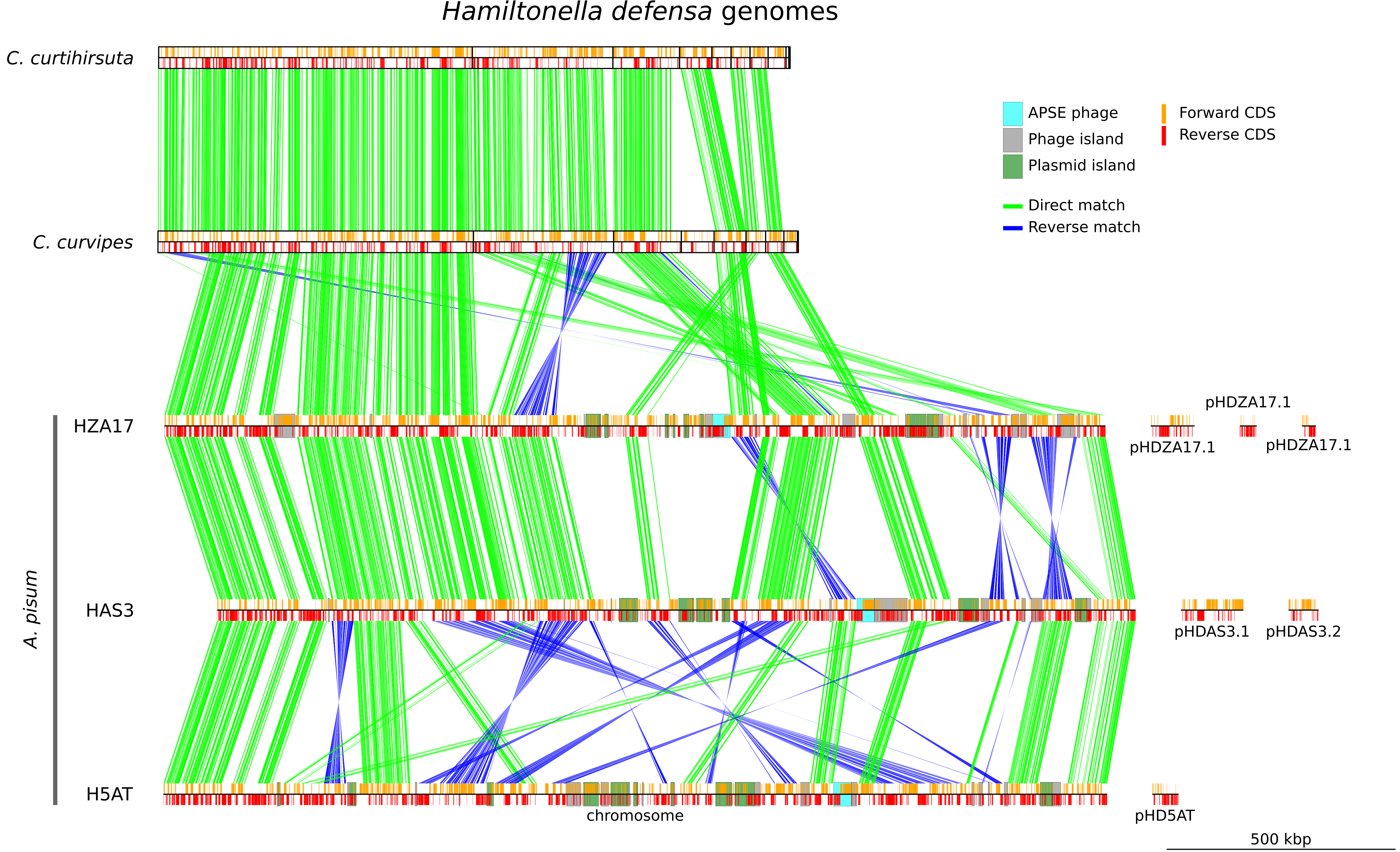


**Fig. S5. Hamiltonella defensa genome synteny and reduction.** Pairwise synteny plots of single-copy core proteins of *Hamiltonella defensa* endosymbionts. *Hamiltonella* symbionts of *Cinara* are plotted with black boxes delimiting the assembled scaffolds. The genomes of facultative *Hamiltonella* symbionts from A. pisum are plotted with their chormosome and plasmids separately, indicating the name of the plasmid either under or over the plotted element. At the bottom right, a scale bar is presented.

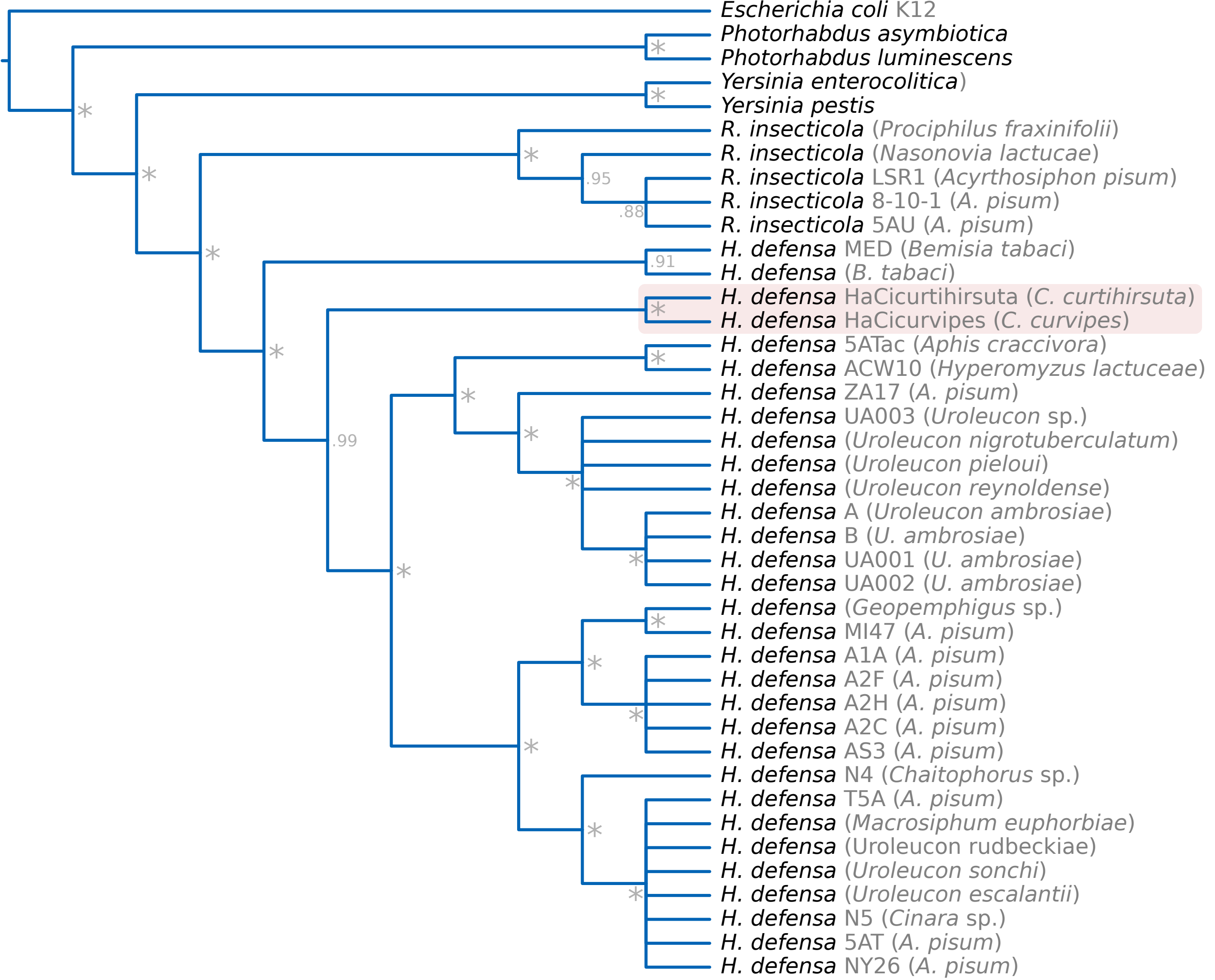


**Fig. S6. Cladogram displaying *Hamiltonella* strain phylogenetic relationships.** Bayesian cladogram showing the relationships among *Hamiltonella* symbiontsinferred from a concatenated alignment of the *accD*, *dnaA*, *gyrB*, *hrpA*, *murE*, *ptsI*, and *recJ* genes. Taxon labels indicate the strain in grey and the host's species in between parenthesis. The monophyletic clade formed by the *Hamiltonella* symbionts from *Cinara* reported in this study is indicated by a red-shaded box.

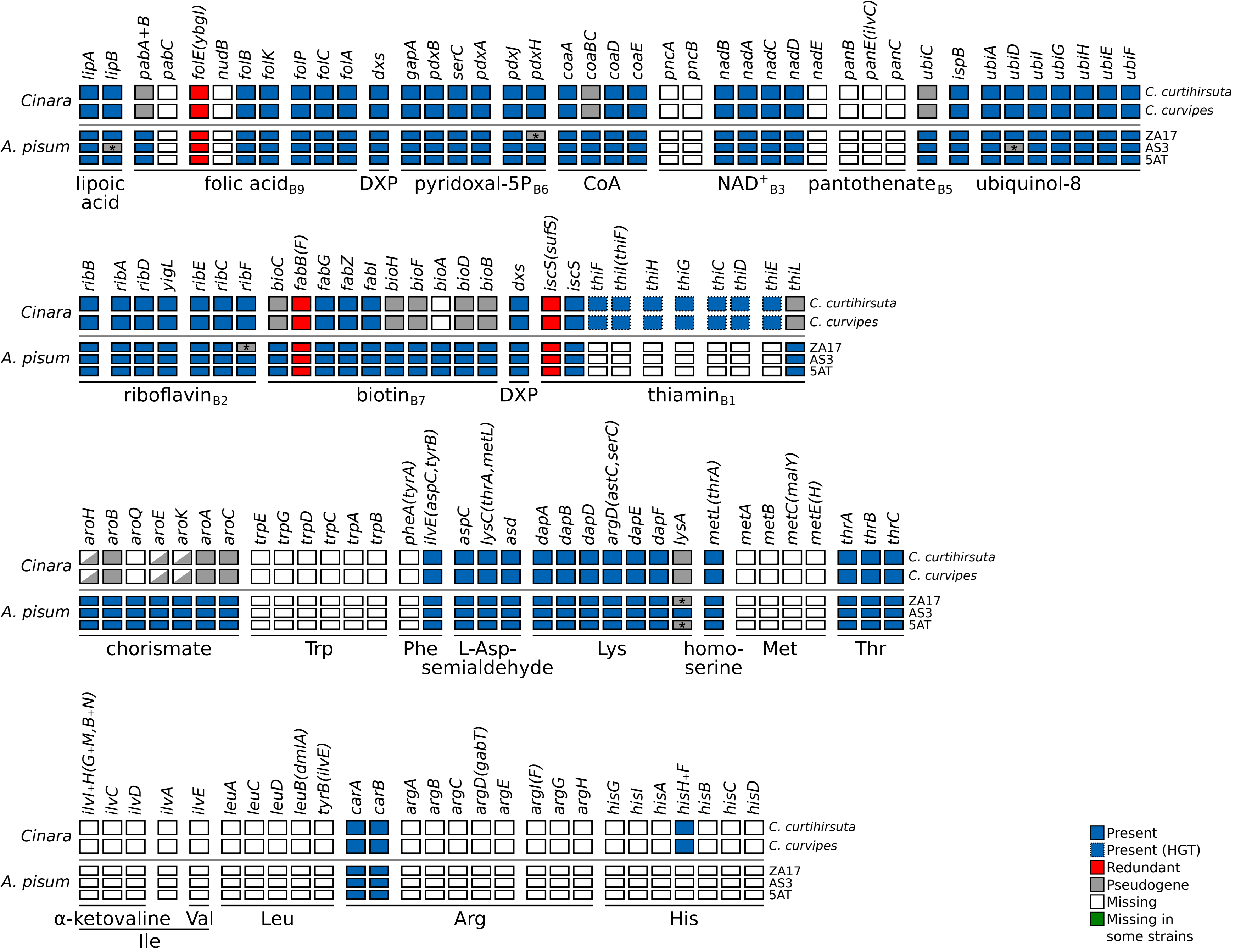


**Fig. S7. Essential B vitamin and amino acid biosynthetic metabolic capabilities of *Hamiltonella defensa* symbionts of different aphid species.** Diagram summarising the metabolic capabilities of the co-obligate *Hamiltonella* symbionts of *Cinara* species and the facultative ones of *A. pisum*. The names of genes coding for enzymes involved in the biosynthetic pathway are used as column names. Each row's boxes represent the genes coded by a symbiont's genome. An asterisk marks a gene pseudogenised by a frameshift in a homopolymeric region. At the right of each row, the host species (for *Cinara* -associated) or symbiont's strain name (for *A. pisum*-associated) of the corresponding symbiont. On the bottom, lines underlining the genes involved in the pathway leading to the compound specified by the name underneath the line.

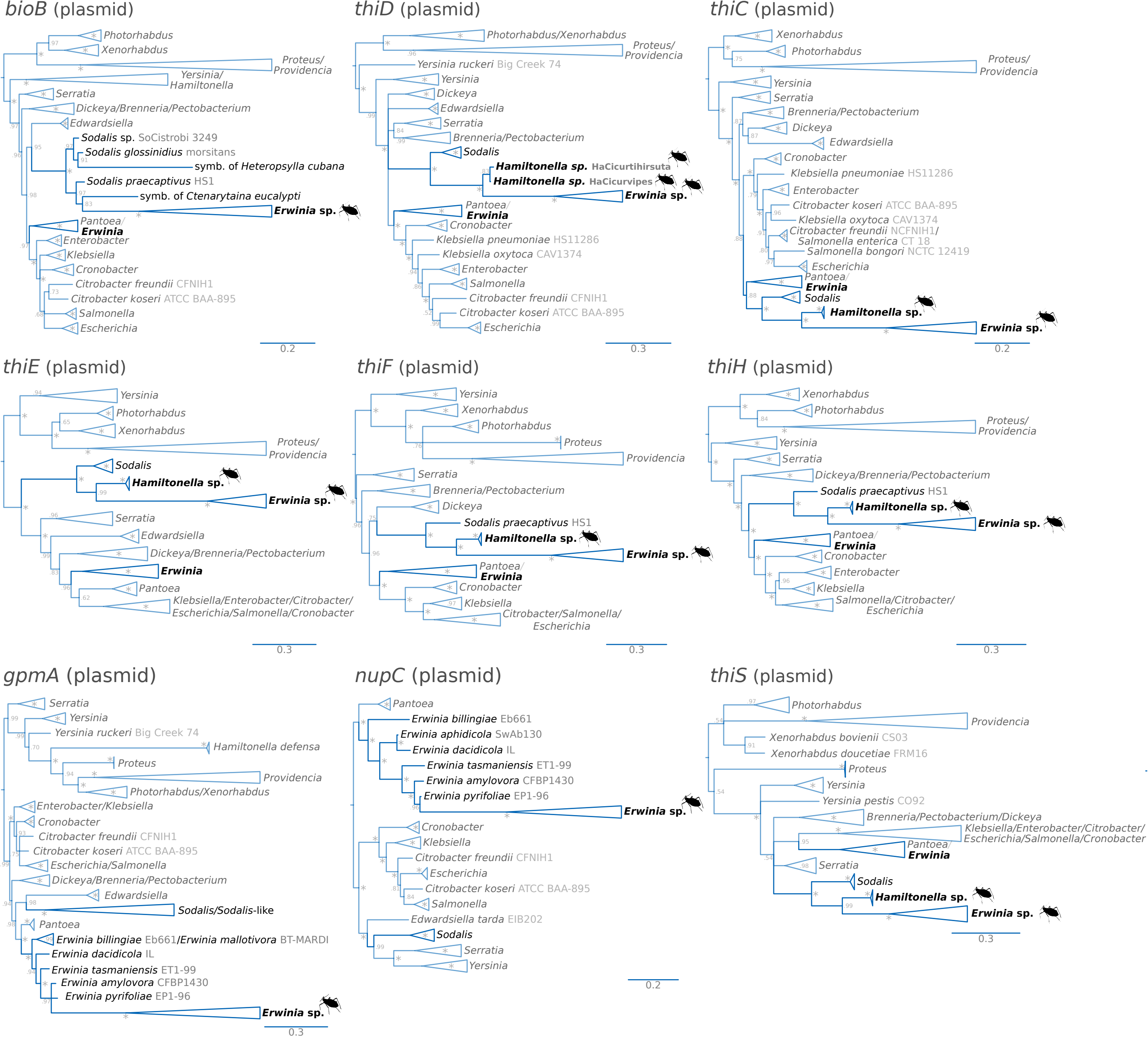


**Fig. S8. Phylograms of biotin- and thiamin-biosynthetic HGT genes and two non-HGT genes (*gpmA* and *nupC*) in plasmids from *Erwinia* symbionts.** Bayesian phylograms of the identified putative HGT (*bioB*, *thiD*, *thiC*, *thiE*, *thiF*, *thiH*, and *thiS*) and non-HGT (*gpmA* and *nupC*) genes present in the plasmids from *Erwinia* symbionts of the newly sequenced *Cinara* species (excluding those from fig. 4). Taxon labels show the strain name in grey. Asterisks at nodes stand for a posterior probability equal to 1. In parenthesis after the gene name, the localisation of the gene in the *Erwinia* genome is indicated. As stated in the main text, whether the *thi* genes in *Hamiltonella* are localised in the chromosome or the a plasmid, remains uncertain.

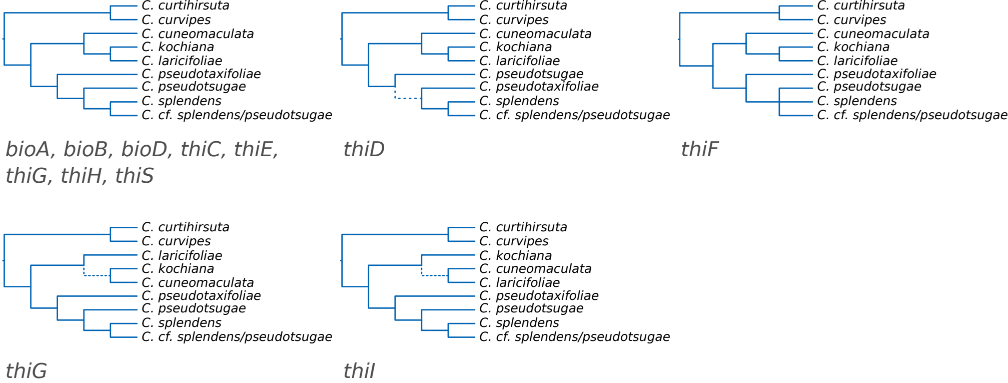


**Fig. S9. Dendograms of the biotin- and thiamin-biosynthetic horizontally-transferred genes.** Dendograms showing the subtree topologies for the biotin- and thiamin-biosynthetic genes originating from a horizontal transfer from *Sodalis*-related bacteria. While most genes show a congruent topology with that of the *Erwinia* and *Buchnera* endosymbionts, the *thiD*, *thiF*, *thiG*, and *thiI* genes have incongruent ones. Dotted branches indicate unsupported (<0.95) bipartitions.

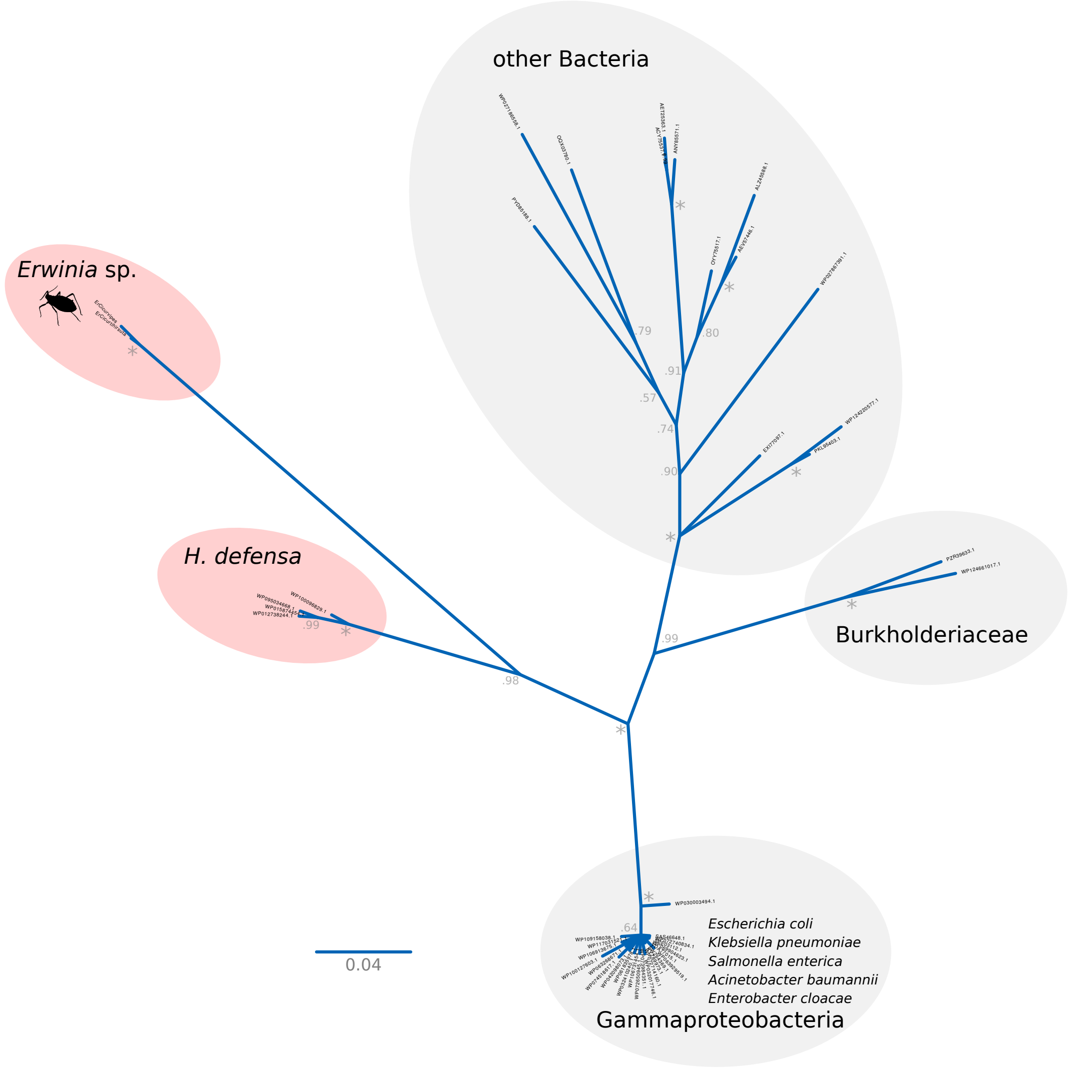


**Fig. S10. Unrooted tree of showing the relationships of Tn3 family resolvase/invertases from *Erwinia* symbionts.** Bayesian tree displaying the well-supported close relationship between Tn3 family resolvase/invertases from *Erwinia* symbionts generated in this study and *Hamiltonella defensa* symbionts.
